# Nuclear Argonaute HRDE-1 Maintains Chromatin-Independent Transcriptional Silencing through a Transgenerational Small RNA Feedback Loop

**DOI:** 10.1101/2025.09.24.678226

**Authors:** Arantxa M. L. Rojas, Almira Chervova, Loan Bourdon, Jagannath Jayaraj, Ana Karina Morao, Olivier Feudjio, Sofija Illic, Sara Capurso, Piergiuseppe Quarato, Julie Ovieve, Germano Cecere

**Affiliations:** Mechanisms of Epigenetic Inheritance, Department of Developmental and Stem Cell Biology, Institut Pasteur, CNRS UMR3738, Paris; Bioinformatics and Biostatistics Hub, Department of Computational Biology, Institut Pasteur, USR 3756, CNRS, Paris; Université Paris Cité, BioSPC, Paris, France

## Abstract

Epigenetic inheritance of transcriptional silencing is traditionally attributed to chromatin-based mechanisms in which Argonaute–small RNA complexes recruit histone-modifying enzymes. Here we show that, in *Caenorhabditis elegans*, the nuclear Argonaute HRDE-1 maintains germline transcriptional repression independently of canonical heterochromatin marks. Using inducible, germline-specific HRDE-1 depletion combined with nuclear sorting, CUT&Tag, and nascent transcription profiling, we identified endogenous genes that become transcriptionally activated upon loss of HRDE-1 despite retaining H3K9me3 and H3K23me3. HRDE-1 directly restrains RNA polymerase II (Pol II) while simultaneously promoting polyUG-dependent amplification of antisense 22G-RNAs in perinuclear condensates, thereby coupling nuclear transcriptional repression to small RNA biogenesis. HRDE-1 loss causes progressive erosion of 22G-RNAs and delays silencing re-establishment, revealing a transgenerational feedback loop in which small RNAs, and not chromatin modifications, constitute the primary heritable signal. These findings redefine nuclear Argonautes as active drivers of RNA-based epigenetic inheritance and broaden our understanding of how small RNA pathways maintain transcriptional silencing across generations.

## Introduction

Epigenetic inheritance of transcriptional silencing is commonly attributed to chromatin-based repression^1^. In this view, nuclear Argonaute proteins bound to small RNAs recruit histone-modifying enzymes to deposit repressive histone marks such as H3K9me3, establishing a chromatin state that can be transmitted across generations^2^. Findings from diverse organisms, from fission yeast to *Drosophila*, have reinforced the idea that canonical heterochromatin is a key determinant of heritable transcriptional silencing. Yet whether such chromatin modifications are universally required for Argonaute-mediated transcriptional silencing remains unresolved.

In *Caenorhabditis elegans*, small RNAs are central to transgenerational epigenetic inheritance^3^. Exogenous double-stranded RNAs (dsRNAs) or endogenous PIWI-interacting RNAs (piRNAs) initiate silencing by triggering the polyUGylation of their RNA targets^4–6^. These modified RNAs serve as templates for RNA-dependent RNA Polymerases (RdRPs) to generate antisense 22-nucleotide small RNAs with a 5′ guanine (22G-RNAs)^5,7,8^. The 22G-RNAs are loaded into a family of Worm-specific Argonautes (WAGOs), which act as silencing effectors^3^. Among them, the nuclear Argonaute HRDE-1 (Heritable RNAi Deficient-1) is required for maintaining transcriptional silencing across generations^6,9,10^. HRDE-1 activity correlates with increased deposition of repressive histone marks, including H3K9me3 and H3K23me3^9,11–13^, reinforcing the long-standing assumption that heterochromatin formation underlies heritable RNA-mediated silencing.

At the same time, heritable RNA silencing can also proceed through distinct, nuclear-independent mechanisms. In some contexts, silencing initiated by exogenous dsRNAs persists in the absence of HRDE-1 or other nuclear factors, relying instead on perinuclear condensates that amplify small RNAs from inherited polyUGylated RNA templates^14,15^. These cytoplasmic amplification systems sustain post-transcriptional gene silencing (PTGS) through cytoplasmic Argonautes and can maintain repression even in the absence of the histone methyltransferases responsible for depositing canonical heterochromatin marks^16^. These observations reveal that heritable silencing can proceed through both nuclear and cytoplasmic routes. Yet the coexistence of these pathways leaves unresolved a central question: when the nuclear pathway is engaged, what molecular features sustain transcriptional repression? In particular, does HRDE-1 require canonical heterochromatin marks to maintain silencing, or can it enforce repression through an alternative, chromatin-independent mechanism?

To address this question, we developed an inducible system for germline-specific HRDE-1 depletion combined with nuclear sorting, CUT&Tag profiling of repressive histone marks^17^, and nascent transcription analysis^18^. This approach enabled direct measurement of transcriptional and chromatin changes at HRDE-1 target loci with germline resolution.

We identify a defined subset of endogenous genes whose silencing depends strictly on HRDE-1 and demonstrate that this transcriptional repression occurs independently of canonical heterochromatin marks. Remarkably, some targets remain decorated with repressive histone modifications yet become transcriptionally active upon loss of HRDE-1, indicating that chromatin marks alone are insufficient to sustain silencing. Moreover, HRDE-1 promotes the transgenerational maintenance of antisense 22G-RNAs through their amplification in perinuclear condensates, thereby coupling nuclear repression to cytoplasmic inheritance.

Together, our findings uncover a chromatin-independent mode of nuclear Argonaute-mediated transcriptional silencing and establish that small RNAs, rather than histone modifications, serve as the primary heritable signal for a subset of germline targets.

## Results

### HRDE-1 is required in every generation to maintain germline gene silencing

Most functional studies of HRDE-1 have relied on transgenic silencing reporters, which, while informative, may not capture the full spectrum of endogenous targets. To systematically define HRDE-1–dependent repression, we combined germline-specific auxin-inducible degradation (AID)^19^ with genome-wide transcriptional profiling (**Fig. 1a** and **Extended Data Fig. 1a**). Acute depletion of HRDE-1 efficiently reduced protein levels (**Extended Data Fig. 1b**) and recapitulated the heritable RNAi defect of the *hrde-1* mutant in a GFP sensor assay (**Extended Data Fig. 1c**). To distinguish immediate effects of HRDE-1 loss during germline development from potential transgenerational consequences, HRDE-1 was depleted for one generation from the first to the last larval stage (G0), and animals were maintained on auxin for four successive generations (G4) (**Fig. 1a** and **Extended Data Fig. 1d,e**). At each generation, synchronized L4-stage worms were collected using a COPAS Biosorter, as previously described^6^, to ensure precise developmental staging across samples.

**Fig. 1:**
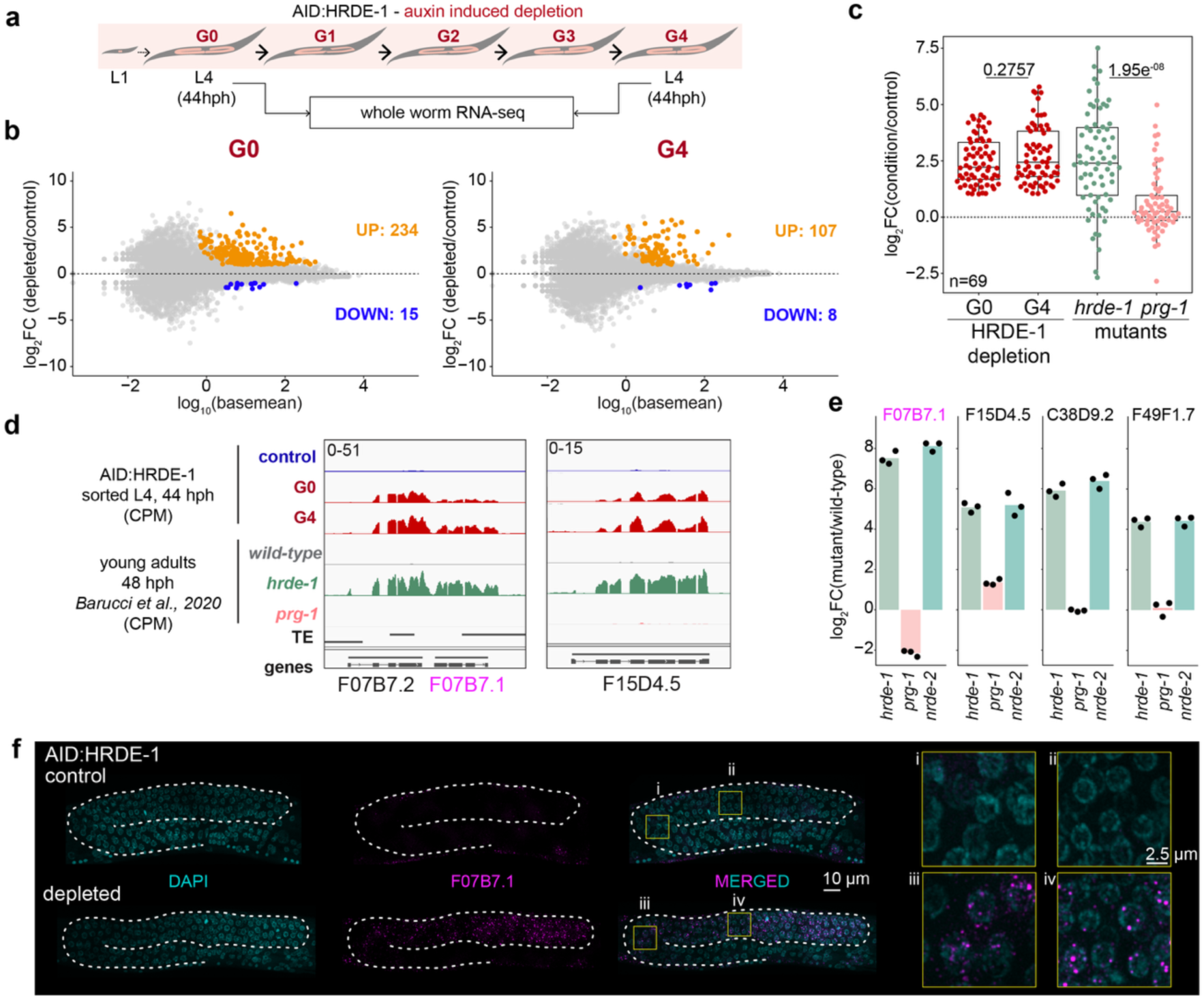
HRDE-1 is required in every generation to maintain germline gene silencing. **a**, Schematic of the worm cultivation strategy for transcriptome profiling. HRDE-1 was depleted in the germline using an auxin-inducible degron from L1 to L4 (G0, 44 hours post-hatching, hph), or continuously for four generations (G4), followed by total RNA-seq from whole worms. **b,** MA plot showing log₂ fold changes (FC) versus mean expression in G0 and G4 animals for protein-coding and pseudogenes. Up-regulated genes (log₂FC ≥ 1, FDR ≤ 0.05) are shown in orange; down-regulated genes (log₂FC ≤ –1, FDR ≤ 0.05) in blue. *n* = 2 biological replicates. **c**, Box plots showing log₂FC for the 69 genes up-regulated in both G0 and G4 (**see Extended Data Fig. 1f**), and their expression changes in *hrde-1* and *prg-1* mutants^20^. P values from Wilcoxon rank-sum tests are indicated. **d,** Genome browser tracks showing normalized RNA-seq coverage (Counts Per Million, CPMs) for representative HRDE-1 targets in control (blue) and HRDE-1 depleted worms (red), and in *hrde-1* (green), *prg-1* (pink) and wild-type controls (grey)^20^. TE, mapped transposable elements. Data show averages of two biological replicates. **e**, RT–qPCR validation of selected HRDE-1–repressed genes in *hrde-1*, *prg-1*, and *nrde-2* mutants at the L4 stage (44 hph), normalized to *act-3*. Bars show mean log₂FC; dots represent individual replicates (*n* = 3). **f**, RNA FISH for F07B7.1 (magenta) in L4 stage HRDE-1–depleted G0 worms compared with controls. Nuclei are marked by DAPI (cyan); gonads are outlined with dashed white lines. Enlarged views of the regions outlined in yellow (**i**, **ii** control; **iii**, **iv** HRDE-1-depleted), are shown alongside each gonad.

Whole-worm RNA-seq revealed that HRDE-1 depletion caused robust de-silencing of endogenous protein-coding genes and pseudogenes in both G0 and G4 animals, with relatively few down-regulated genes (**Fig. 1b**). These results indicate that most transcriptional changes reflect direct HRDE-1–mediated repression. As expected, the total number of mis-regulated genes was lower than in the *hrde-1* mutant^20^, consistent with incomplete depletion using the degron system. Importantly, the number of up-regulated genes did not increase from G0 to G4 (**Fig. 1c and Extended Data Fig. 1f**), showing that HRDE-1 must act in every generation to maintain germline silencing rather than functioning through a purely transgenerational mechanism.

To distinguish HRDE-1–specific targets from those regulated by piRNAs, we compared the 69 genes de-silenced in both G0 and G4 (**Extended Data Fig. 1f**) with published RNA-seq datasets from *hrde-1 (tm1200)* and *prg-1 (n4357)* mutants^20^. Most of these genes were up-regulated in *hrde-1* but not in *prg-1* mutants (**Fig. 1c,d**). RT-qPCR confirmed that loss of PRG-1 had minimal effect on their expression (**Fig. 1e**). In contrast, mutation of *nrde-2*, a nuclear silencing cofactor^9,21^, resulted in de-silencing comparable to *hrde-1* (**Fig. 1e**). These results demonstrate that these loci are bona fide targets of the nuclear WAGO pathway (hereafter referred to as HRDE-1–repressed targets).

Because these genes remain repressed independently of PRG-1, they are distinct from the spermatogenic, piRNA-dependent 22G-RNA targets previously characterized^6^. Consistently, RNA FISH^22^ for one of the most strongly de-silenced genes, F07B7.1, showed robust expression throughout the germline of HRDE-1–depleted animals but not in controls (**Fig. 1f** and **Extended Data Fig. 1g**). This expression pattern differs from piRNA-regulated spermatogenic genes, which are transiently expressed in pachytene nuclei^6^. Together, these observations indicate that HRDE-1–repressed targets represent a distinct class of constitutively silenced germline loci that provide a tractable system to dissect the molecular mechanisms of HRDE-1–mediated transcriptional repression and small-RNA inheritance.

### HRDE-1 depletion in G0 causes transcriptional de-silencing in germline nuclei

To determine whether HRDE-1–dependent repression occurs directly within the germline nucleus, we isolated germline nuclei from control and HRDE-1–depleted worms using a GFP nuclear reporter and fluorescence-activated nuclei sorting (**Fig. 2a** and **Extended Data Fig. 2a-c**). RNA-seq from sorted nuclei quantified steady-state nuclear RNA (**Fig. 2b**), and GRO-seq measured nascent transcription (**Fig. 2c**). Both datasets revealed similar numbers of up-regulated genes and very few down-regulated ones (**Fig. 2b,c**), mirroring the whole-worm RNA-seq results (**Fig. 1**). The concordant increase detected by both RNA-seq and GRO-seq indicates that HRDE-1 represses transcription directly within germline nuclei, rather than acting predominantly through post-transcriptional mechanisms (**Fig. 2d**).

**Fig. 2:**
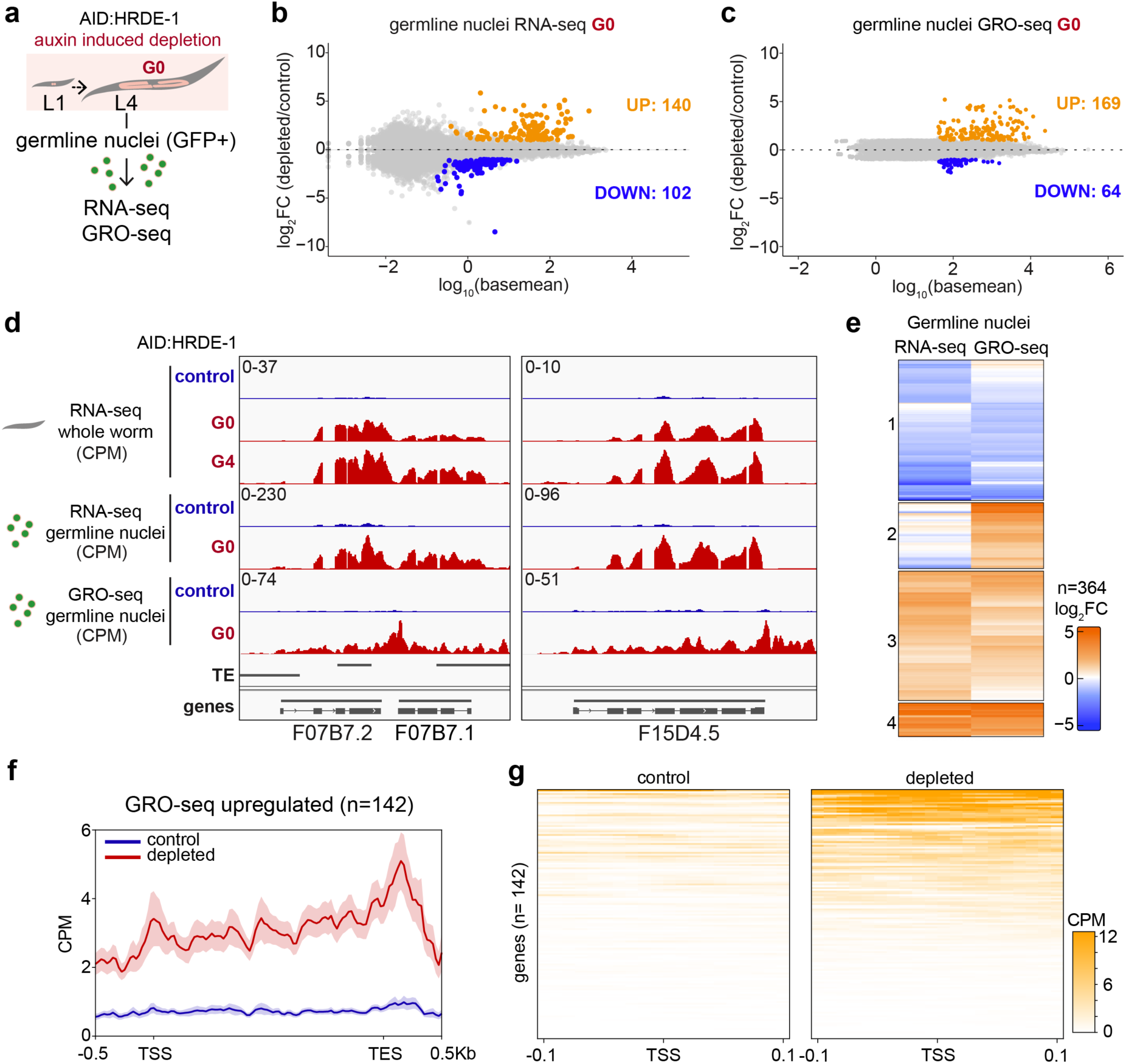
HRDE-1 depletion in G0 causes transcriptional de-silencing in germline nuclei. **a,** Schematic of germline nuclei isolation after one generation of HRDE-1 depletion (G0) for RNA-seq and GRO-seq analysis (**see Extended Data Fig. 2a-c**). **b, c,** MA plots showing log₂ fold changes (FC) versus mean expression for protein-coding and pseudogenes in RNA-seq (**b**) and GRO-seq (**c**) from G0 germline nuclei. Up-regulated genes (log₂FC ≥ 1, FDR ≤ 0.05) are shown in orange; down-regulated genes (log₂FC ≤ –1, FDR ≤ 0.05) in blue. *n* = 4 biological replicates. **d,** Genome browser views of representative targets, showing normalized RNA-seq and GRO-seq read coverage (Counts Per Million, CPM) in control (blue) and HRDE-1-depleted (red) germline nuclei. For comparison RNA-seq coverage from whole-worm G0 and G4 samples (as in **Fig. 1d**) is shown. TE, mapped transposable elements. Data represent averages from two (whole worm) or four (sorted nuclei) biological replicates. **e,** Heatmap summarizing significant expression changes (FDR ≤ 0.05) for genes de-silenced in G0 HRDE-1–depleted germline nuclei in either RNA-seq or GRO-seq. **f,** Metagene profiles of GRO-seq read coverage for the 142 genes significantly up-regulated in GRO-seq, shown for control (blue) and HRDE-1-depleted (red) L4 worms. TSS, transcription start site; TES, transcription end site. **g,** Heatmap showing GRO-seq read coverage around the TSS (±100 bp) for the 142 genes analyzed in **(f)**, illustrating widespread increases in nascent transcription upon HRDE-1 depletion.

Across the 364 genes exhibiting significant expression changes (FDR ≤ 0.05) in either dataset, most showed comparable fold changes in both assays (**Fig. 2e**), confirming that these represent bona fide transcriptional targets of HRDE-1.

To characterize how HRDE-1 influences transcription dynamics, we examined nascent RNA profiles for the 142 genes significantly up-regulated in GRO-seq. We excluded an additional 27 genes whose transcription start sites (TSSs) were located <500 bp upstream of an expressed gene, where read-through transcription from neighboring loci could confound the signal interpretation. Metagene analysis across gene bodies and within ±500 bp of the TSS and transcription end sites (TES) revealed elevated GRO-seq signal along the entire gene body, with pronounced peaks at both TSS and TES in HRDE-1–depleted samples (**Fig. 2f**). Notably, upstream of the TSS, transcriptional activity also remained higher than in controls, indicating enhanced transcription initiation. A heatmap of GRO-seq signal within ±100 bp of TSSs confirmed that this increase was pervasive across nearly all affected genes (**Fig. 2g**).

These findings demonstrate that HRDE-1 loss causes a broad increase in nascent transcription, including at initiation sites, challenging the prevailing view that nuclear Argonautes repress transcription primarily by inhibiting elongation. This contrasts with the somatic nuclear Argonaute NRDE-3, which blocks Pol II during elongation^21^, suggesting that nuclear Argonautes employ distinct regulatory mechanisms in different tissues. Altogether, acute depletion of HRDE-1 from L1 to L4 is sufficient to transcriptionally de-silence its germline targets, revealing a germline-specific mode of nuclear Argonaute-mediated transcriptional repression.

### HRDE-1 maintains silencing independently of canonical heterochromatin marks

Silencing by nuclear Argonautes is generally associated with the deposition of repressive histone modifications^2,9,11–13^. To test whether the transcriptional de-repression observed upon HRDE-1 depletion results from changes in these marks, we developed a germline-specific CUT&Tag approach that combines GFP-based nuclear sorting with histone modification profiling, enabling high-resolution chromatin analysis in the germline (**Fig. 3a, Extended Data Figs. 2-5**, and **Methods**). This strategy addresses two major limitations of previous studies: whole-worm profiling, which dilutes germline signals with abundant somatic H3K9me3; and the widespread use of an anti-H3K9me3 antibody (Abcam 8898) recently shown to cross-react with H3K27me3 and H4K20me3 in *C. elegans*^23^. We validated an alternative antibody (Active Motif 39161) with high specificity for H3K9me3 (**Extended Data Fig. 3a-c**). As a functional test, germline-specific depletion of SET-25^24^—the sole H3K9me3 methyltransferase—between L1 and L4 completely erased H3K9me3 peaks in sorted nuclei (**Extended Data Fig. 3d,e**), confirming both antibody specificity and the sensitivity of our approach.

**Fig. 3:**
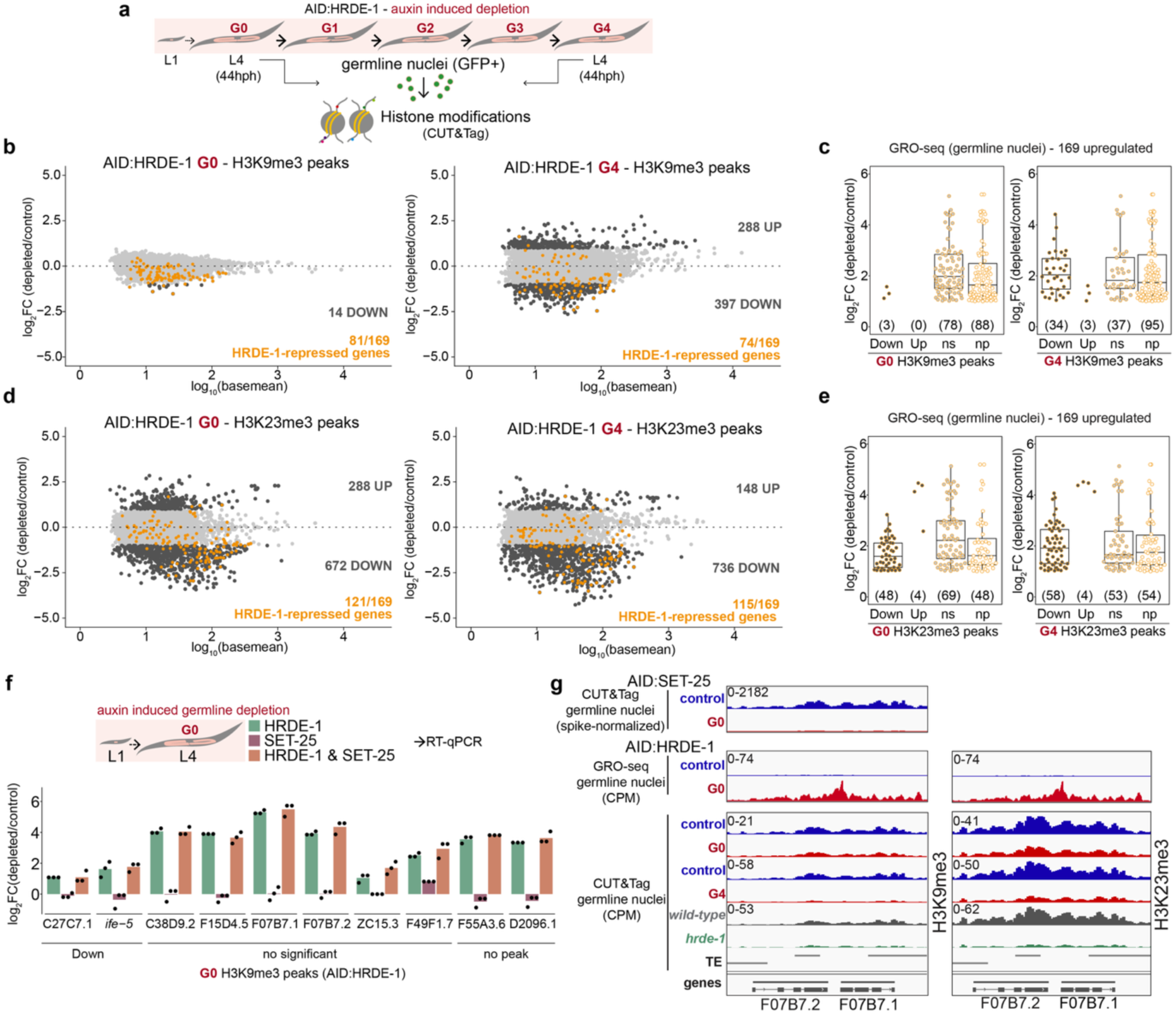
HRDE-1 maintains transcriptional silencing independently of canonical heterochromatin marks. **a**, Schematic of germline nuclei isolation after HRDE-1 depletion for one generation (G0) or four generations (G4) for histone modification profiling by CUT&Tag. **b**, MA plots of H3K9me3 CUT&Tag signal in germline nuclei of G0 and G4 HRDE-1–depleted worms. Genes up-regulated in G0 GRO-seq (from **Fig. 2c**) that intersect H3K9me3 peaks are highlighted in orange; significantly altered H3K9me3 peaks (log₂FC ≥ 1 or ≤ –1, FDR ≤ 0.05) are shown in dark grey. *n* = 4 biological replicates. **c**, Box plots showing log₂ fold change (FC) in H3K9me3 CUT&Tag signal for the 169 G0-upregulated genes (Fig. 2c) that intersect H3K9me3 peaks. Categories: *ns* (no significant change with peak present), *np* (no peak detected), *Down* (significantly decreased H3K9me3), and *Up* (significantly increased H3K9me3). The number of genes in each category is indicated in parentheses. **d**, MA-plots of H3K23me3 CUT&Tag signal in germline nuclei from G0 and G4 HRDE-1–depleted worms, with color coding as in (**b**). *n* = 4 biological replicates. **e**, Box plots of log₂FC for up-regulated genes (Fig. 2c) intersecting with H3K23me3 peaks. Categories as in (**c**). **f**, RT–qPCR validation of selected HRDE-1–repressed genes in germlines depleted of HRDE-1 (green), SET-25 (pink), or both HRDE-1 and SET-25 (dark orange), normalized to *act-3*. Bars show mean log₂FC; dots represent individual replicates (*n* = 3). The category for each gene (as in **c**) is indicated below. **g,** Genome browser views of representative targets, showing H3K9me3 and H3K23me3 CUT&Tag coverage (Counts Per Million, CPM) in G0 and G4 HRDE-1–depleted germline nuclei (control, blue; HRDE-1 depletion, red), as well as wild-type (grey) and *hrde-1* mutant (green). H3K9me3 CUT&Tag signal in G0 SET-25–depleted germline nuclei (G0; spike-in normalized counts, see **Extended Data Fig. 3d,e**) is shown in comparison. GRO-seq signal from G0 HRDE-1–depleted nuclei (Fig. 2d) is included to indicate transcriptional up-regulation. TE, mapped transposable elements. Data are averaged from three or four biological replicates.

We next examined H3K9me3 at genes showing transcriptional changes in GRO-seq after one generation (G0) of HRDE-1 or SET-25 depletion (**Fig. 2c**, **Fig. 3a-c,g,** and **Extended Data Fig. 4a-e**). Of 169 up-regulated genes, 81 and 89 overlapped with H3K9me3 peaks, yet 78 showed no significant reduction in H3K9me3 signal upon HRDE-1 depletion (log₂FC ≤ 1, FDR ≤ 0.05) (**Fig. 3b,c**). Only three genes exhibited both loss of H3K9me3 and transcriptional up-regulation (**Fig. 3c**). Conversely, most down-regulated genes displayed unchanged H3K9me3 levels (**Extended Data Fig. 4d,e**). Depletion of SET-25 alone did not trigger de-silencing, and simultaneous loss of HRDE-1 and SET-25 did not enhance transcriptional activation regardless of H3K9me3 status (**Fig. 3f**). These results show that loss of H3K9me3 is insufficient to explain transcriptional activation and support a model in which SET-25 acts downstream of HRDE-1–mediated repression.

To assess transgenerational effects, we profiled H3K9me3 after four generations of HRDE-1 depletion (G4) and in *hrde-1* knockout worms (**Fig. 3b,c,g**, and **Extended Data Fig. 4f-m**). Both conditions showed a larger number of altered peaks, including both gains and losses, indicating widespread chromatin remodeling in the absence of HRDE-1. The number of up-regulated genes overlapping H3K9me3 peaks remained similar (**Fig. 3b right, Extended Data Fig. 4l**), but genes with decreased H3K9me3 peaks increased from 3 in G0 to ∼33–34 in G4 and *hrde-1* knockout animals (**Fig. 3c right, Extended Data Fig. 4m**). This expansion reflects progressive chromatin remodeling that accumulates across generations, rather than a primary cause of transcriptional de-silencing. Notably, several HRDE-1–repressed genes lacked detectable H3K9me3 or retained wild-type levels even when transcriptionally activated, further demonstrating that SET-25–dependent H3K9me3 deposition occurs downstream of HRDE-1 and is not required for repression.

We next analyzed H3K23me3, another repressive modification implicated in HRDE-1–mediated silencing^16^. One generation of HRDE-1 depletion caused broader changes in this mark than in H3K9me3, with 288 peaks increased and 672 decreased (**Fig. 3d,g** and **Extended Data Fig. 5a**). Among the 169 transcriptionally up-regulated genes, 121 overlapped H3K23me3 peaks (**Fig. 3d**), yet most (69) showed no significant change, 48 decreased, and 4 increased. Thus, H3K23me3 remodeling is largely uncoupled from transcriptional outcome (**Fig. 3d,e,g**, and **Extended Data Fig. 5b,c**). Moreover, increased H3K23me3 did not correlate with repression (**Extended Data Fig. 5b,c**). Extended HRDE-1 depletion (G4) or loss of *hrde-1* increased the amplitude but not the diversity of H3K23me3 changes (**Fig. 3d right,g**, and **Extended Data Fig. 5d-k**), again consistent with secondary chromatin remodeling rather than a primary regulatory role.

We also considered whether HRDE-1–dependent silencing involves changes in the active promoter mark H3K4me3^25^. Germline-specific CUT&Tag profiling of H3K4me3 after one generation of HRDE-1 depletion showed only modest genome-wide changes, with 30 peaks gained. Among the 169 up-regulated genes, only 12 displayed increased H3K4me3 signal (**Extended Data Fig. 5l–o**), and most de-silenced genes lacked detectable H3K4me3 enrichment. Thus, transcriptional activation of HRDE-1 targets cannot be explained by the accumulation or redistribution of this active mark.

Collectively, these results show that HRDE-1 maintains transcriptional silencing in the germline independently of canonical heterochromatin modifications. Transcriptional de-repression occurs even when H3K9me3 and H3K23me3 remain at wild-type levels, and depletion of the H3K9 methyltransferase does not phenocopy HRDE-1 loss. We conclude that repressive histone mark deposition, the accompanying chromatin remodeling, and changes in active promoter marks are downstream consequences of HRDE-1–mediated transcriptional silencing rather than its cause.

### HRDE-1 promotes the transgenerational maintenance of 22G-RNAs

Having established that HRDE-1 sustains transcriptional silencing independently of canonical heterochromatin marks, we next investigated whether its function is linked to maintaining the small RNA populations required for transgenerational silencing. Loss of HRDE-1 has previously been associated with a large-scale reduction in antisense 22G-RNAs^20^, and artificial tethering of HRDE-1 can trigger *de novo* production of heritable small RNAs^26^, suggesting a direct role in their biogenesis.

We sequenced small RNAs from G0, G1, G2, and G4 HRDE-1–depleted L4 worms (**Fig. 4a,b** and **Extended Data Fig. 6a**). Across generations, antisense 22G-RNAs progressively decreased, with the extent of dysregulation increasing in each successive generation (**Fig. 4b**). This cumulative loss cannot be explained solely by reduced stability of existing small RNAs and instead reflects a transgenerational requirement for HRDE-1 in promoting 22G-RNA biogenesis.

**Fig. 4:**
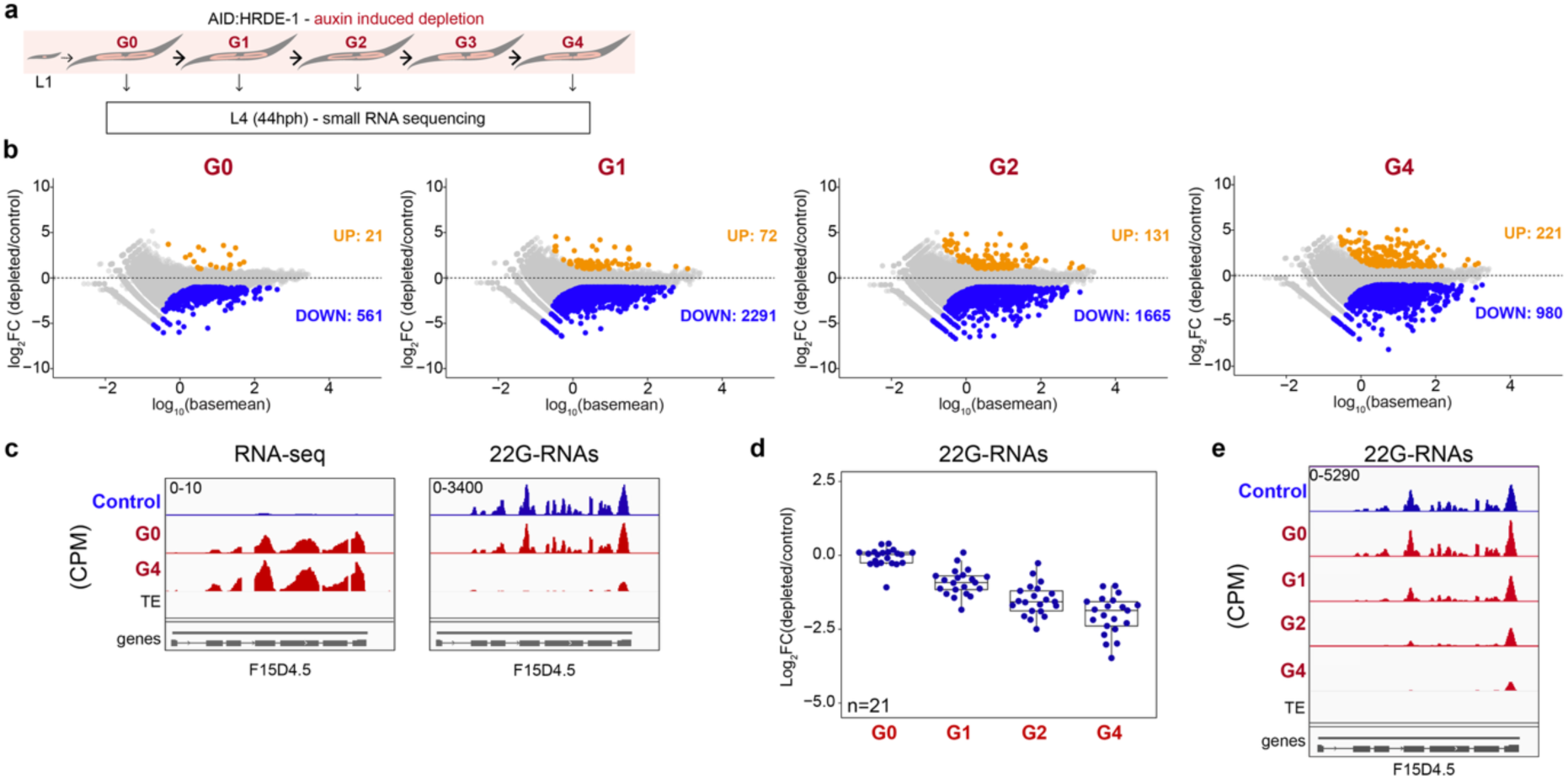
HRDE-1 is required for the transgenerational maintenance of 22G-RNAs. **a,** Experimental design for small RNA-seq across generations following germline-specific HRDE-1 depletion. HRDE-1 was depleted during development (G0) or continuously for one (G1), two (G2), or four (G4) generations. **b**, MA plot showing 22G-RNAs mapping antisense to protein-coding genes and pseudogenes in HRDE-1–depleted versus control animals. Orange indicates significantly increased 22G-RNAs (log₂FC ≥ 1, FDR ≤ 0.05); blue indicates significantly decreased 22G-RNAs (log₂FC ≤ –1, FDR ≤ 0.05). *n* = 2 biological replicates. **c**, Genome browser views of normalized RNA-seq (left) and 22G-RNA (right) coverage (Counts Per Million, CPM) for F15D4.5, a representative HRDE-1–repressed gene. mRNA expression is similarly de-silenced in G0 and G4 (RNA-seq), whereas 22G-RNAs are markedly reduced only in G4. Control, blue; HRDE-1–depleted, red. TE, mapped transposable elements. *n* = 2 biological replicates. **d**, Box plot showing log₂ fold change (FC) in 22G-RNAs at G0, G1, G2, and G4 for the 21 HRDE-1–repressed genes that lose small RNAs in G4 (**see Extended Data Fig. 6d**). **f**, Genome browser views illustrating the gradual, generation-dependent reduction of 22G-RNAs antisense to F15D4.5 in G0, G1, G2, and G4. Control, blue; HRDE-1–depleted, red. TE, mapped transposable elements. *n* = 2 biological replicates.

Although 22G-RNAs were also reduced in G0 animals, none of the 69 genes consistently de-silenced upon HRDE-1 depletion (**Extended Data Fig. 1f**) showed decreased 22G-RNAs at this early generation (**Fig. 4c** and **Extended Data Fig. 6b,c**). Thus, transcriptional de-repression in G0 occurs without detectable changes in small RNAs, and conversely, small RNA loss does not necessarily coincide with transcriptional activation (**Fig. 4c** and **Extended Data Fig. 6b,c**). This indicates that the 22G-RNAs available for loading into cytoplasmic WAGOs are insufficient to maintain repression post-transcriptionally when HRDE-1 is absent, contrasting with previous observations in RNAi inheritance, where post-transcriptional silencing can compensate for the loss of nuclear HRDE-1^14^. Our approach, therefore, identifies a subset of endogenous loci that rely exclusively on HRDE-1-mediated transcriptional silencing, revealing a nuclear mode of heritable repression that cytoplasmic WAGOs cannot buffer.

The number of genes losing 22G-RNAs increased markedly across generations (**Fig. 4a,b** and **Extended Data Fig. 6b**), whereas the number of de-silenced genes remained constant (**Fig. 1a,b**). By G4, 21 of the 69 de-silenced genes had significantly reduced antisense 22G-RNAs in both replicate experiments (**Extended Data Fig. 6c**). These genes displayed no reduction in G0 but a pronounced and progressive loss of 22G-RNAs by G4, with a graded decline across generations (**Fig. 4d,e**).

Collectively, these data indicate that HRDE-1 has two separable roles: it directly enforces transcriptional silencing in the germline nucleus and promotes the transgenerational maintenance of 22G-RNA populations through small RNA amplification.

### Mutator foci, but not Z granules, are required for HRDE-1–mediated repression

We next asked whether HRDE-1–dependent 22G-RNA amplification requires cytoplasmic perinuclear condensates or occurs independently of condensates within the nucleus. To define the cytoplasmic requirements of HRDE-1–dependent small-RNA maintenance, we first assessed whether HRDE-1 targets require polyUGylation^5^, the modification that primes mRNAs for 22G-RNA synthesis and occurs primarily within Mutator foci condensates (**Fig. 5a-c,e**). PolyUGylation was strongly impaired in *hrde-1* mutants, comparable to *rde-3* mutants lacking the polyUG ribonucleotidyltransferase (**Fig. 5c,e**), indicating that HRDE-1 is required for this step of small RNA amplification. In contrast, polyUGylation of piRNA-dependent targets, such as D2096.1 and F55A6.3, was unaffected in *hrde-1* mutants, consistent with previous findings^6^ (**Fig. 5c**). These results demonstrate that the HRDE-1–repressed genes identified here form a distinct class of piRNA-independent targets, whose silencing and small RNA amplification rely on HRDE-1 activity but not on a piRNA trigger.

**Fig. 5:**
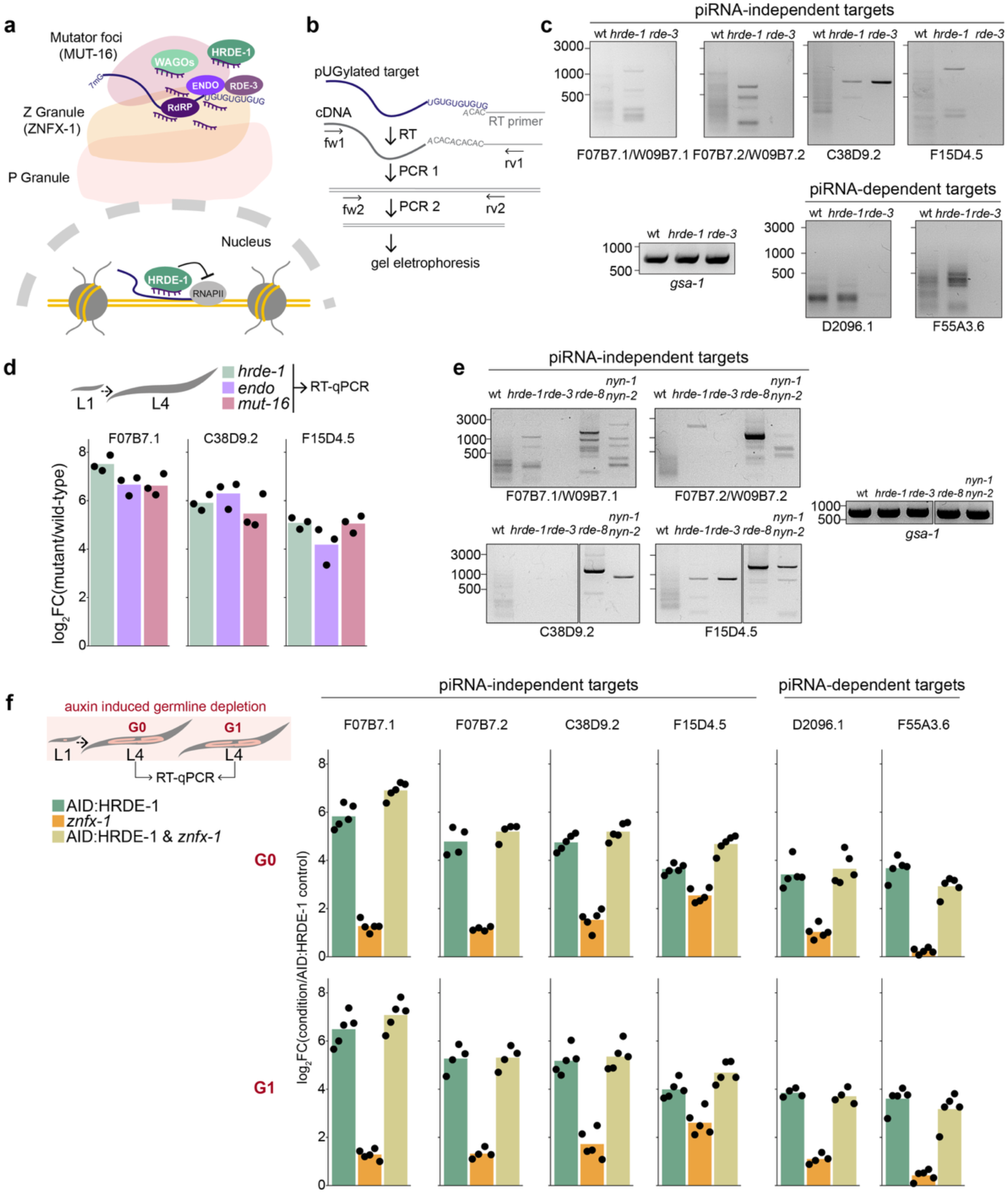
Mutator foci, but not Z granules, are required for HRDE-1–mediated repression. **a**, Schematic of the perinuclear condensates present in *C. elegans* germlines, where 22G-RNA biogenesis occurs, highlighting the main components analyzed in this study. ENDO, endonucleases (RDE-8, NYN-1, NYN-2); RdRP, RNA-dependent RNA polymerase. **b**, Schematic of polyUGylation assay, based on oligo-specific reverse transcription (RT primer shown in grey) of polyUGylated mRNAs and PCR amplification using reverse primers complementary to adaptor sequences in the RT primer (rv1 and rv2) and gene-specific primers (fw1 and fw2). **c,e,** PolyUGylation assay for HRDE-1–repressed genes that lose 22G-RNAs transgenerationally, analyzed across different genetic backgrounds. *gsa-1* serves as a control. Genes are grouped by their dependence on the piRNA pathway for 22G-RNA biogenesis^20^. **d**, RT–qPCR analysis of HRDE-1–repressed genes in *hrde-1*, *rde-8;nyn-1;nyn-2* triple mutants (“endo”), and *mut-16* mutants at the L4 stage (44 hours post-hatching, hph). Expression values are normalized to *act-3*. Bars show mean log₂ fold change (FC), dots represent biological replicates (*n* = 3). **f**, RT–qPCR analysis of selected HRDE-1–repressed genes in control and *znfx-1* mutant backgrounds following G0 or G1 HRDE-1 depletion at the L4 stage, normalized to *act-3*. Bars show mean log₂FC; dots represent biological replicates (*n* = 4-5). Genes are grouped as in (**c**).

We next examined the endonucleases RDE-8, NYN-1, and NYN-2, which generate RNA fragments for polyUGylation^5,27^. PolyUGylation defects in *hrde-1* mutants mirrored those observed in *rde-8* mutants or *nyn-1;nyn-2* double mutants (**Fig. 5e**). Correspondingly, expression of HRDE-1 targets in *rde-8;nyn-1;nyn-2* triple mutants (“endo”) was comparable to *hrde-1* mutants (**Fig. 5d**).

Because these enzymes localize to Mutator foci, we asked whether these condensates are required for HRDE-1–dependent transcriptional repression. Loss of MUT-16, the scaffold protein that nucleates Mutator foci^28^, phenocopied *hrde-1* mutants and caused similar de-silencing of HRDE-1 targets (**Fig. 5d**). These results demonstrate that Mutator foci are essential for both 22G-RNA amplification and nuclear repression of HRDE-1-regulated genes, serving as the cytoplasmic interface through which HRDE-1 couples nuclear target recognition to small RNA biogenesis.

Given the established role of Z granules in post-transcriptional RNAi inheritance^14,29^, we next asked whether they also contribute to HRDE-1–mediated transcriptional silencing. Z granules, nucleated by the helicase ZNFX-1^29^, support RNAi inheritance by amplifying small RNAs through a nuclear-independent mechanism^14,15^. Publicly available small RNA-seq dataset^14^ and our polyUGylation assays showed reduced 22G-RNAs and impaired polyUGylation for several HRDE-1 targets in *znfx-1* mutants (**Extended Data Fig. 7a–c**), similar to *hrde-1* mutants. However, RT-qPCR, and RNA FISH revealed only mild or no transcriptional de-repression in *znfx-1* mutants relative to wild type (**Fig. 5f** and **Extended Figure 7d,e**). Moreover, depletion of HRDE-1 in a *znfx-1* mutant background did not enhance de-repression in either G0 or G1 (**Fig. 5f**).

Together, these findings show that although Z granules influence small RNA abundance, they are dispensable for maintaining transcriptional silencing of HRDE-1 targets. Instead, HRDE-1-mediated repression relies on Mutator foci, which couple nuclear transcriptional silencing with heritable small RNA amplification and thereby establish a mechanistic bridge between the nuclear and cytoplasmic arms of the inheritance pathway.

### Silencing recovery mediated by HRDE-1 requires 22G-RNAs

By acutely depleting HRDE-1, we uncovered the generational dynamics of silencing maintenance. We next investigated how repression is re-established once HRDE-1 expression is restored. Worms depleted of HRDE-1 for four generations (G4) were removed from auxin treatment and propagated for one (R1), four (R4), and six (R6) generations before collection at the L4 stage (**Fig. 6a** and **Extended Data Fig. 1a,d,e**). Notably, while HRDE-1 depletion during germline development (G0) was sufficient to de-silence its targets (**Fig. 1** and **Fig. 6b,c**), re-expressing HRDE-1 during the same developmental period did not restore repression in R1 (**Fig. 6b,c**). RNA-seq revealed that the 21 HRDE-1-repressed genes requiring HRDE-1 for small RNA maintenance remained strongly up-regulated in R1 (**Fig. 6b**). Silencing was gradually re-established over subsequent generations, with some genes still de-silenced in R6 (**Fig. 6b,c**).

**Fig. 6:**
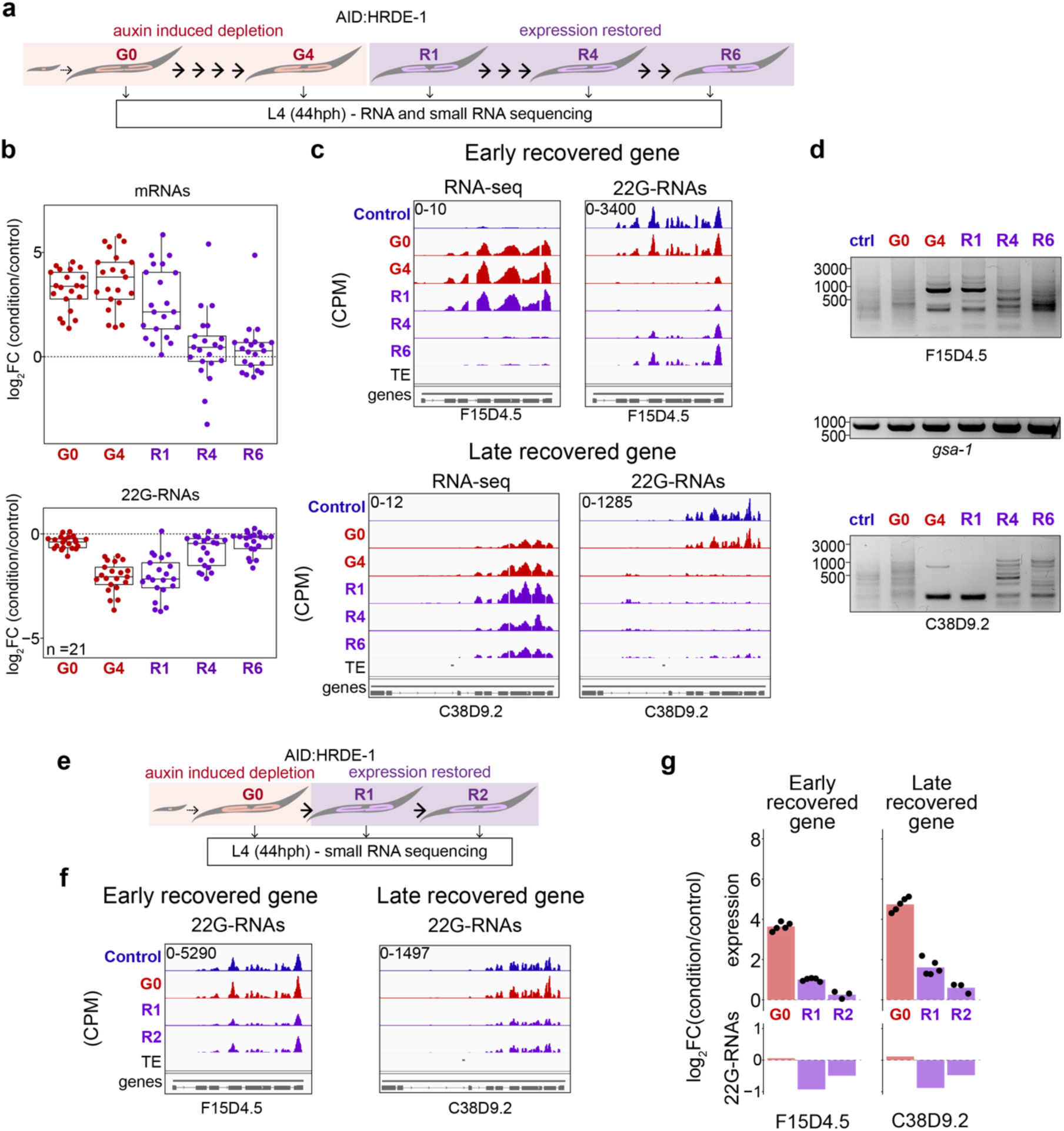
Maintenance and re-establishment of HRDE-1–dependent silencing requires 22G-RNA biogenesis. **a**, Experimental design to assess transgenerational maintenance and recovery of HRDE-1–dependent silencing. **b**, Box plots showing log₂ fold change (FC) of mRNA levels (top) and 22G-RNAs (bottom) for the 21 HRDE-1–repressed genes that lose small RNAs in G4 (as in **Fig. 4d**). Values are shown for G0 and G4 HRDE-1–depleted worms (red) and for R1, R4, and R6 worms after HRDE-1 re-expression (purple). **c**, Genome browser views of normalized RNA-seq and 22G-RNAs coverage (Counts Per Millions, CPM) for two representative HRDE-1–repressed genes with distinct recovery kinetics: F15D4.5 (early recovered) and C38D9.2 (late recovered). Control samples are shown in blue, HRDE-1-depleted samples in red, and recovery samples in purple. TE, mapped transposable elements. **d**, PolyUGylation assay for F15D4.5 and C38D9.2 across control (blue), G0 and G4 HRDE-1 depletion (red), and R1, R4, and R6 recovery generations (purple). *gsa-1* serves as a control. **e**, Experimental design testing silencing recovery after only one generation (G0) of HRDE-1 depletion, followed by R1 and R2 recovery. **f,** Genome browser views of normalized 22G-RNA coverage (CPM) for F15D4.5 (early recovered) and C38D9.2 (late recovered) in G0 (red), R1 and R2 (purple), and control (blue) samples. **g**, RT–qPCR analysis of F15D4.5 and C38D9.2 expression in G0, R1, and R2 worms at the L4 stage (44 hours post-hatching, hph), normalized to *act-3*. Bars show mean log_2_FC; dots represent biological replicates (*n* = 3–5). Bottom: corresponding average FC of 22G-RNAs for each gene across generations.

We classified genes as “early recovered” if both transcriptional repression and 22G-RNA levels were restored by R4, and as “late recovered” if they remained de-silenced and showed incomplete 22G-RNA recovery even by R6. This classification highlights substantial gene-specific variability in HRDE-1–dependent silencing re-establishment and indicates that restoration of repression requires multiple rounds of germline transmission.

Because most targets exhibited a pronounced loss of antisense 22G-RNAs by G4, we next measured 22G-RNA levels during recovery. In R1, failure to restore repression correlated with a complete absence of 22G-RNA recovery, indicating that HRDE-1 requires these small RNAs to re-engage transcriptional silencing (**Fig. 6b,c**). In later generations, 22G-RNAs progressively re-accumulated in parallel with transcriptional repression, consistent with a model in which small RNA regeneration, like their maintenance, is transgenerationally promoted by HRDE-1.

To determine whether recovery depends on the same cytoplasmic biogenesis steps required for maintenance, we monitored polyUGylated RNA intermediates for two HRDE-1 targets with distinct recovery kinetics, F15D4.5 (early recovered) and C38D9.2 (late recovered) (**Fig. 6c,d**), alongside with the piRNA-dependent target *bath-13*, whose 22G-RNA biogenesis is HRDE-1–independent^6^ (**Extended Data Fig. 8a**). PolyUGylation of *bath-13* remained stable across generations, whereas F15D4.5 and C38D9.2 showed a strong loss at G4 and no recovery in R1 (**Fig. 6c,d** and **Extended Data Fig. 8a**). PolyUG signal increased by R4 and R6. However, for C38D9.2, the pattern remained weaker and more discrete than in controls, matching its slower transcriptional repression. These findings suggest that gene-specific differences in how efficiently HRDE-1 promotes polyUGylation underlie variable recovery kinetics, echoing earlier observations^16^.

Finally, to directly test whether recovery failure in R1 results from the absence of 22G-RNAs, we repeated the experiment using only one generation of HRDE-1 depletion (G0) (**Fig. 6e**). Because 22G-RNA levels remain intact in G0 for de-silenced genes, we predicted that restoring HRDE-1 in the next generation would enable immediate repression. Indeed, small RNA sequencing in R1 and R2 after G0 depletion confirmed wild-type 22G-RNA levels (**Fig. 6e,f**), and RT-qPCR showed that repression was fully re-established in R1 (**Fig. 6g** and **Extended Data Fig. 8b**). Together, these findings demonstrate that silencing recovery by HRDE-1 requires pre-existing antisense 22G-RNAs. Once these small RNAs are lost, their biogenesis must be rebuilt gradually across generations, establishing a requirement for a transgenerational small RNA feedback loop in the re-establishment of HRDE-1–mediated transcriptional repression.

## Discussion

For nearly two decades, the prevailing model has held that nuclear Argonautes repress transcription by recruiting histone methyltransferases to establish repressive chromatin. In *C. elegans*, genetic links between HRDE-1, the NRDE cofactors, and the H3K9 methyltransferases SET-25 and SET-32 have consistently positioned heterochromatin at the core of heritable gene silencing^9,16,30^. By acutely depleting HRDE-1 specifically in the germline and combining this with transcriptional and chromatin profiling of sorted germline nuclei, we now identify a class of endogenous loci whose repression depends on HRDE-1 yet persists independently of canonical heterochromatin marks. These findings redefine the relationship between nuclear Argonautes and chromatin modifiers. HRDE-1 maintains transcriptional silencing without requiring histone methylation, indicating that Argonaute-mediated control of Pol II can occur independently of heterochromatin and that chromatin remodeling may follow, rather than drive, transcriptional repression.

Our germline-resolved CUT&Tag and GRO-seq analyses show that HRDE-1 loss increases Pol II occupancy and nascent transcription across entire gene bodies, including promoter regions, despite largely intact H3K9me3 or H3K23me3. Depletion of SET-25 alone does not cause transcriptional de-repression, and chromatin changes accumulate only after multiple generations without HRDE-1, consistent with a secondary consequence of persistent transcription. Together, these results demonstrate that canonical heterochromatin marks are not strictly required for HRDE-1-mediated silencing. Supporting this model, previous proteomic analyses detected Pol II subunits, but not histone methyltransferases, among HRDE-1 interactors^31^. HRDE-1, therefore, likely represses transcription directly at the level of Pol II recruitment or initiation. This mechanism differs from that of the somatic nuclear Argonaute NRDE-3, which inhibits Pol II elongation^21^, highlighting tissue-specific specialization among nuclear Argonautes.

Comparable chromatin-independent repression has been reported in *Drosophila*, where the Piwi–piRNA pathway can initiate transcriptional repression independently of HP1a and H3K9me3^32–34^. However, these studies relied on forced Piwi tethering to synthetic reporters and primarily captured the initiation step, as stable silencing of endogenous transposons still required heterochromatin. In contrast, HRDE-1 acts at endogenous loci to maintain and transmit transcriptional repression across generations without heterochromatin involvement. By coupling direct nuclear inhibition of Pol II to continual small RNA amplification in cytoplasmic condensates, HRDE-1 establishes a self-sustaining feedback loop that reinstates repression each generation. Thus, chromatin-independent repression by nuclear Argonautes is not limited to initiation but can support long-term, heritable silencing.

The HRDE-1–repressed genes identified here represent a distinct class of constitutively silenced germline loci, clearly separable from the piRNA-triggered spermatogenic targets governed by PRG-1^6^. Spermatogenic targets undergo cyclic, developmentally programmed expression followed by repression in each generation, driven by piRNA-guided initiation of 22G-RNA production. In contrast, HRDE-1–dependent loci remain constitutively silenced throughout germline development and across generations, maintaining repression without renewed piRNA input. Several of these loci overlap with “predicted influencers of RNA-regulated expression” (PIREs) and include F07B7.2/SDG-1, recently characterized by the José laboratory^35,36^, highlighting their recurrent sensitivity to multiple RNA-silencing pathways.

HRDE-1–dependent 22G-RNA biogenesis requires the endonucleases RDE-8 and NYN-1/2, which cleave target RNAs upstream of polyUGylation by RDE-3, and an intact Mutator-foci scaffold provided by MUT-16. In contrast, Z granules, which mediate post-transcriptional inheritance, are dispensable for nuclear repression. Although the Z-granule helicase ZNFX-1 reduces 22G-RNA abundance at these loci, ZNFX-1 does not enhance silencing defects. These findings indicate that the relevant small RNA amplification for transcriptional repression occurs predominantly in Mutator foci, arguing against the notion of a nuclear small RNA amplification mechanism^14,26^, and establishing HRDE-1 as the central factor linking nuclear target recognition to cytoplasmic small RNA production.

Our depletion–recovery experiments further illuminate how this coupling enforces transgenerational memory. When HRDE-1 is absent for multiple generations, silencing recovery proceeds heterogeneously across targets. Genes such as F15D4.5 recover transcriptional silencing and 22G-RNA levels by the fourth generation (“early-recovered”), whereas others like C38D9.2 remain de-repressed and exhibit incomplete 22G-RNA restoration after six generations (“late-recovered”). These distinct kinetics likely reflect differences in HRDE-1’s efficiency at promoting polyUGylation and in the accessibility of polyUGylated templates for small RNA amplification. Thus, recovery requires HRDE-1 to re-engage both nuclear repression and small RNA regeneration, creating a feedback loop in which each process reinforces the other until the heritable silenced state is restored.

Together, these findings refine the molecular logic of epigenetic inheritance. While chromatin modifications clearly contribute to the long-term repression, they are not the decisive regulatory input for a specific subset of HRDE-1-regulated genes. Instead, small RNAs themselves constitute the primary heritable signal that maintains transcriptional states across generations. By decoupling transcriptional silencing from heterochromatin yet linking it to small RNA amplification in mutator foci, HRDE-1 defines a self-reinforcing feedback circuit that bridges nuclear transcriptional control with cytoplasmic small RNA biogenesis. This RNA-centered inheritance mechanism unifies initiation, maintenance, and transmission of silencing into a single framework and suggests that analogous feedback loops between transcription and small RNA pathways may underlie adaptive gene regulation in other biological contexts.

## Extended Data Figures

**Extended Data Fig. 1:**
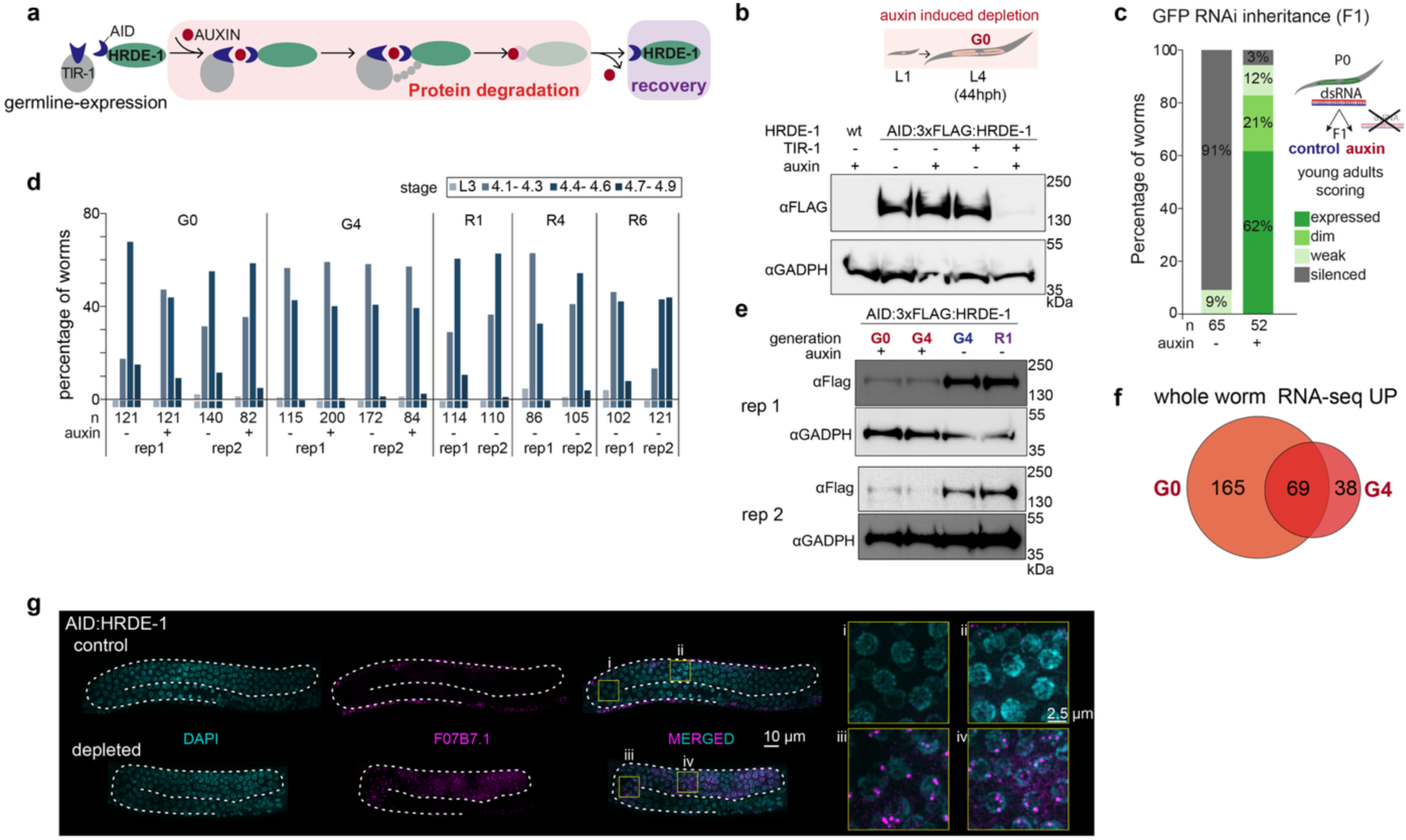
Validation of the auxin-inducible degron system and experimental setup for HRDE-1 depletion and recovery. **a**, Schematic of the auxin-inducible degron (AID) strategy used for germline-specific HRDE-1 depletion. **b**, Western blot of AID:3xFLAG:HRDE-1 in wild-type (untagged HRDE-1) and AID-tagged strains with or without germline TIR-1 expression, in the presence or absence of auxin. Anti-FLAG antibody was used for HRDE-1 detection; GAPDH served as a loading control. **c**, RNAi inheritance assay in AID::3xFLAG::HRDE-1; GFP::piRNA control sensor worms injected with dsRNA against GFP and grown on ethanol (control) or auxin (HRDE-1 depleted). Silencing was assessed in the F1 generation. **d**, Stage distribution of L4 larvae sorted by COPAS Biosorter for RNA-seq and small RNA-seq (two biological replicates) used in **Fig. 1a,b, Fig. 6a-d**, and **Extended Data Fig. 6b-d**. **e**, Western blot of L4 larvae at G0 and G4 (HRDE-1–depleted, red) and R1 (recovery, purple), compared to control (blue). Two biological replicates corresponding to sequencing samples are shown. **f**, Venn diagram showing the intersection between G0 and G4 up-regulated genes (log₂FC ≥ 1, FDR ≤ 0.05) identified by RNA-seq (Fig. 1b). **g,** RNA FISH for *F07B7.1* (magenta) of G4 HRDE-1–depleted L4 larvae compared to control. DAPI (cyan) marks nuclei; gonads are outlined with dashed white. Insets show magnified regions (i, ii = control; iii, iv = HRDE-1–depleted).

**Extended Data Fig. 2:**
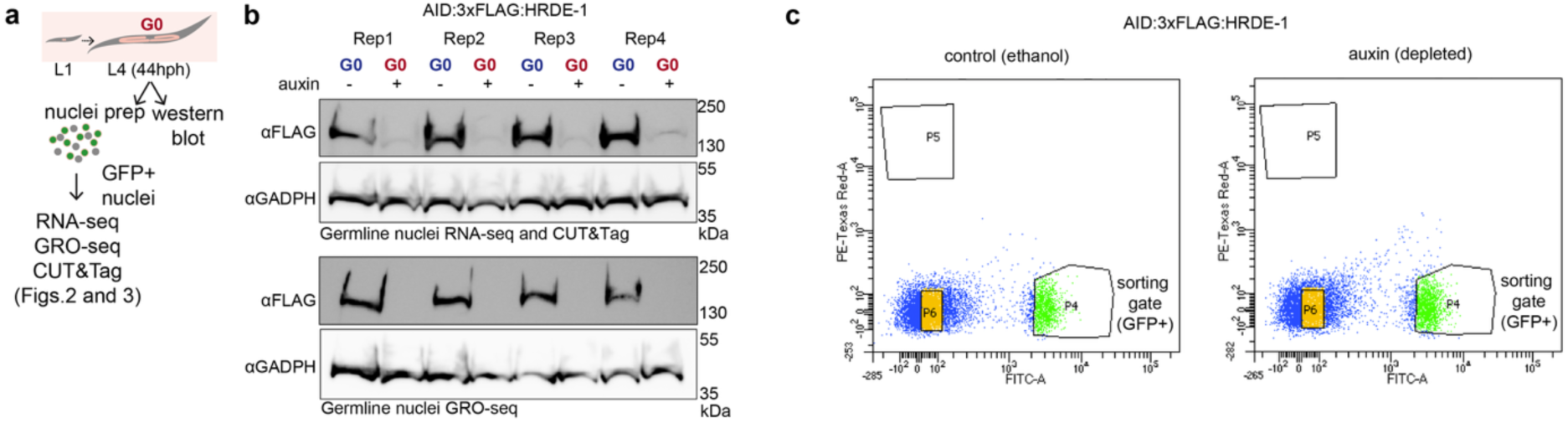
Germline nuclei isolation workflow for HRDE-1–depleted worms. **a,** Schematic of germline nuclei isolation from HRDE-1–depleted worms for downstream analyses (Corresponding to **Figs. 2 and 3**). **b,** Western blot of L4 lysates used for **Fig. 2 and Fig. 3** confirming HRDE-1 depletion in each replicate (*n* = 4 biological replicates). **c,** Representative FACS plots for nuclei sorting. X-axis: FITC-A (GFP detection); Y-axis: PE-Texas Red-A (no expected events). P4 = GFP-positive nuclei; P6 = non-GFP events; P5 = no expected events.

**Extended Data Fig. 3:**
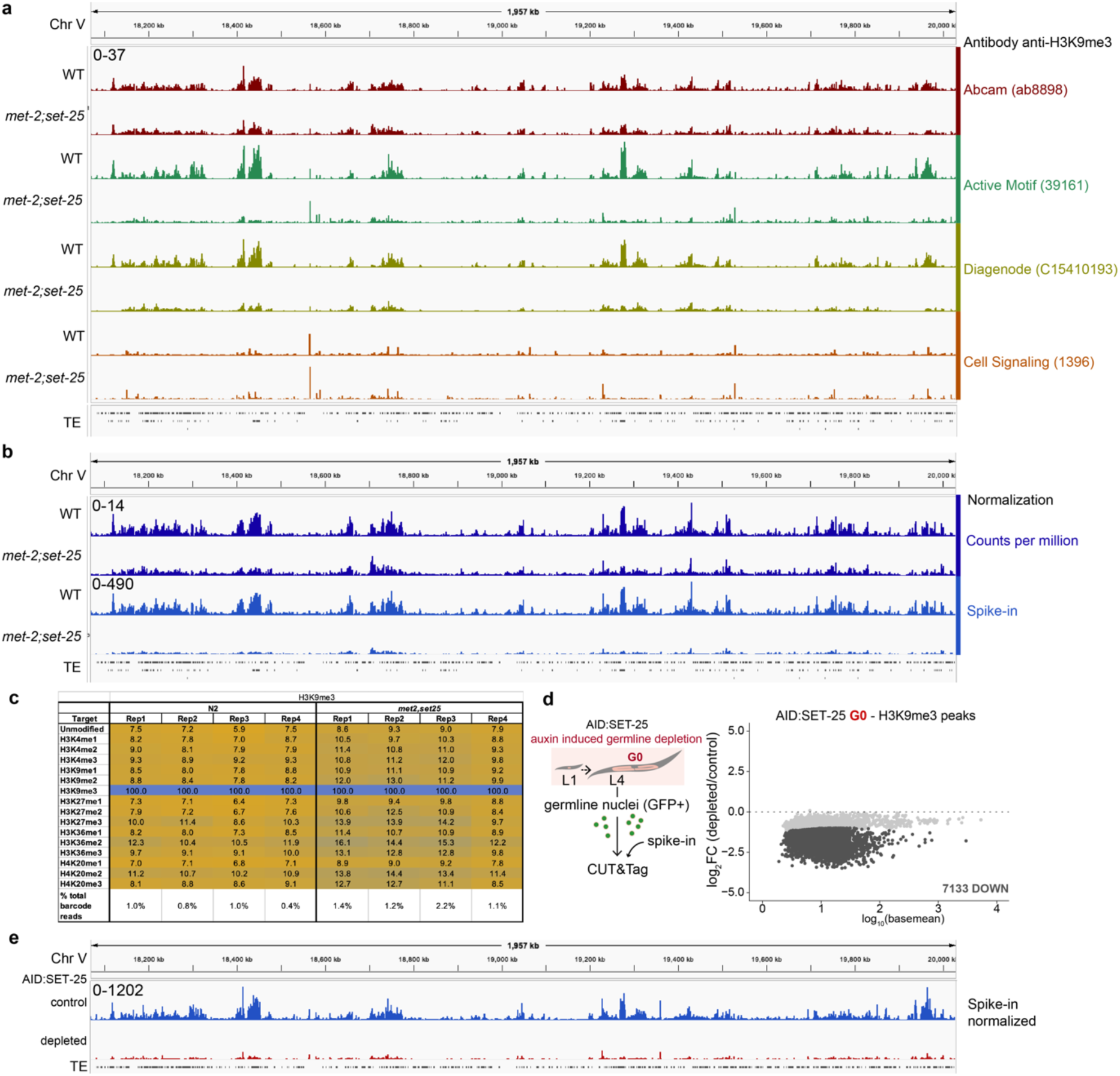
CUT&Tag strategy and antibody validation for H3K9me3 profiling in sorted germline nuclei. **a,** Genome browser views of a representative region on Chromosome V, showing normalized CUT&Tag signal (Count Per Million, CPM) using four antibodies (right) to detect H3K9me3 in wild-type and *met-2;set-25* L4 worms. **b,** CUT&Tag tracks of the same region as in **a**, using the Active Motif H3K9me3 antibody with synthetic nucleosomes spike-in (Epicypher) in wild-type and *met-2;set-25* sorted L4 nuclei. Coverage is normalized either by total reads (CPM) or spike-in scaling. **c,** Antibody specificity table (Epicypher) for the H3K9me3 antibody used in **b**. **d,** Schematic of germline-specific SET-25 depletion and sample preparation for CUT&Tag. Right: MA plot showing H3K9me3 changes upon germline SET-25 depletion (G0); significantly changed peaks are shown in dark grey. **e,** Genome browser views of the region shown in **a**, comparing CUT&Tag signal in G0 SET-25-depleted germline nuclei and controls. Coverage is normalized using spike-in-derived scaling factors. TE,mapped transposable elements.

**Extended Data Fig. 4:**
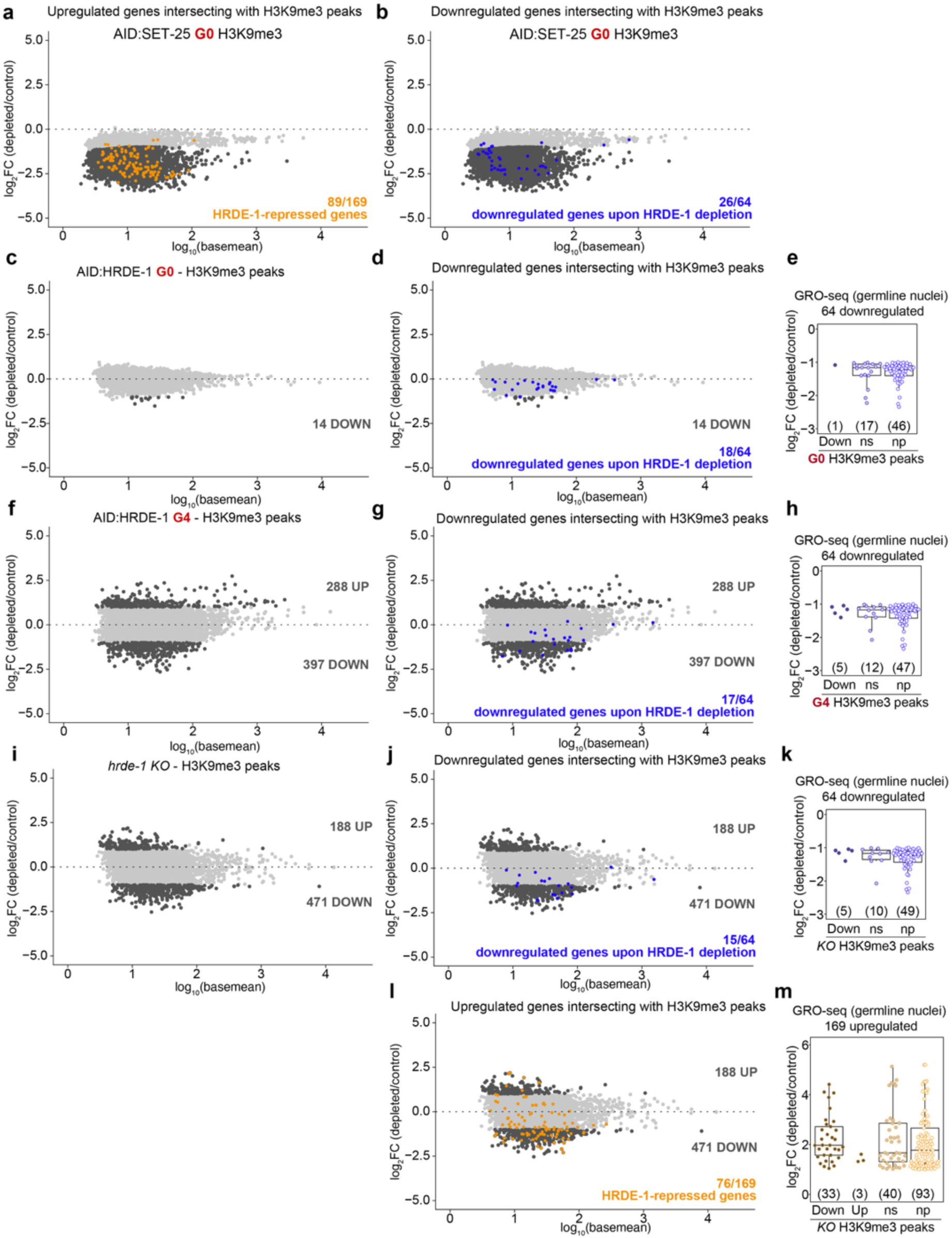
HRDE-1 maintains transcriptional silencing independently of H3K9me3. **a**, MA plot of H3K9me3 CUT&Tag signal upon SET-25 depletion. Genes up-regulated in HRDE-1–depleted nuclei (**Fig. 2c**) that intersect H3K9me3 peaks are highlighted in orange. **b**, Same as **a**, but highlighting genes down-regulated in HRDE-1–depleted nuclei (blue). **c**,**f,i**, MA plots of H3K9me3 signal after one generation of HRDE-1 depletion (G0) **(c)**, four generations (G4) **(f),** or in *hrde-1* knockout germline nuclei (KO) **(i)**. Significantly changed peaks are shown in dark grey (log₂FC ≥ 1 or ≤ –1, FDR ≤ 0.05). **d**,**g,j,l,** Same dataset as **(c),(f),(i**), highlighting intersection with down-regulated genes (blue; **c, f, i**) or up-regulated genes (orange; **l**) from **Fig. 2c**. **e**,**h,k,m,** Box plots of log₂ fold change (FC) for down-regulated (**e,h,k**) or up-regulated (**m**) genes grouped by H3K9me3 status: *ns* (no significant change, peak present), *np* (no peak), Down (significantly reduced peak), or Up (significantly increased peak). Category counts indicated in parentheses.

**Extended Data Fig. 5:**
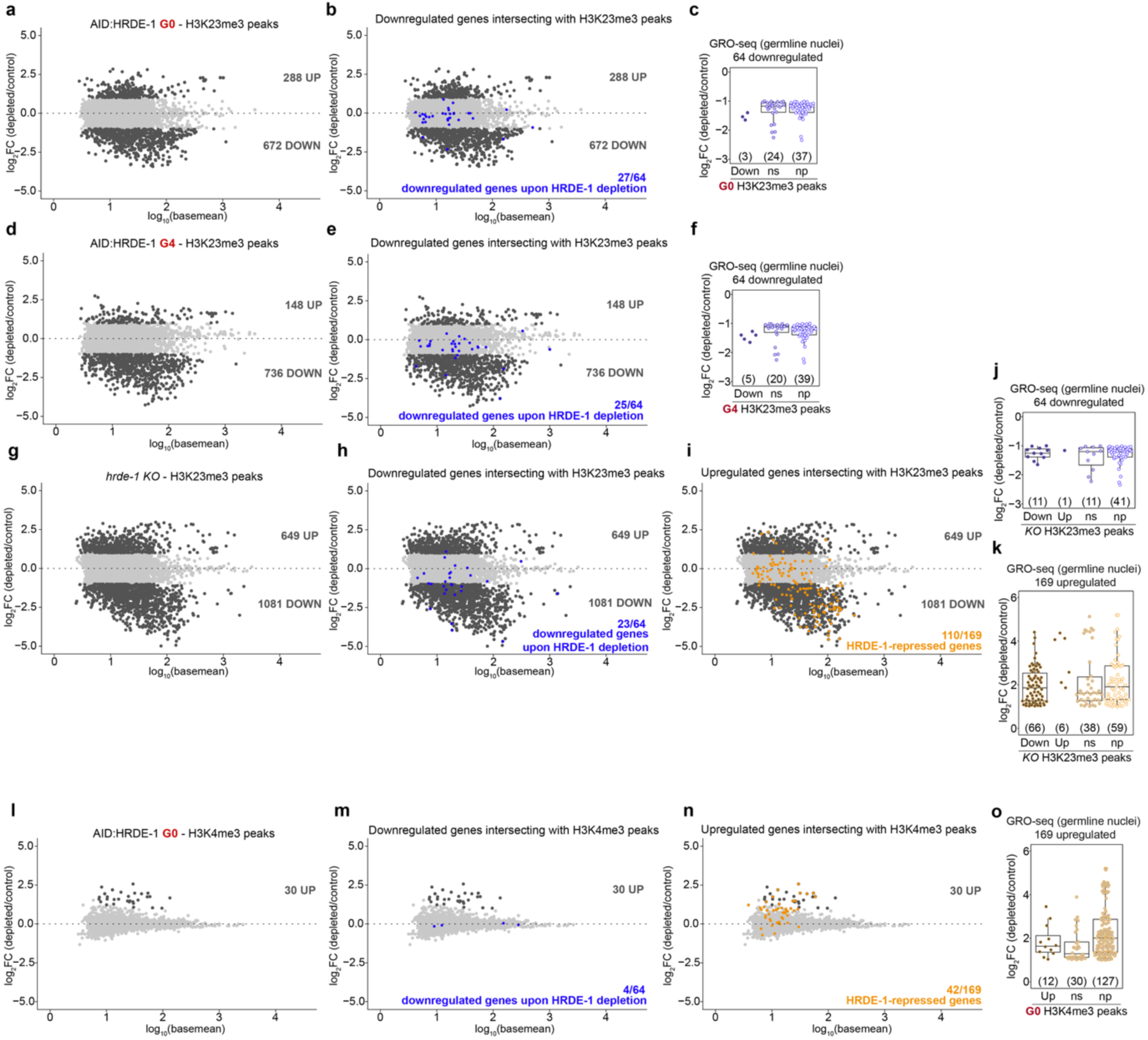
HRDE-1–dependent silencing occurs independently of changes in H3K23me3 or H3K4me3. **a,d,g,** MA plots of H3K23me3 CUT&Tag signal upon HRDE-1 depletion for one generation (G0) (**a**), four generations (G4) (**d**), or in *hrde-1* knockout germline nuclei (KO) (**g**). Significantly changed peaks are in dark grey (log₂FC ≥ 1 or ≤ –1, FDR ≤ 0.05). **b**,**e,h,** Same as in **a**,**d,g**, but highlighting intersection with down-regulated (blue) or up-regulated (orange) from **Fig. 2c**. **c**,**f,j,k,** Box plots of log₂ fold change (FC) for down-regulated (**c,f,g**) or up-regulated (**k**) genes grouped by H3K23me3 status: *ns* (no significant change, peak present), *np* (no peak), Down (significantly reduced peak), or Up (significantly increased peak). Category counts indicated in parentheses. **l**, MA plot of H3K4me3 signal upon HRDE-1 depletion (G0). **m,n**, Highlighting intersections between (**l)** and down-regulated (**m**, blue) or up-regulated (**n**, orange) genes from **Fig. 2c**. **o,** Box plot of log₂FC for up-regulated genes grouped by H3K4me3 status: *ns*, *np*, Up.

**Extended Data Fig. 6:**
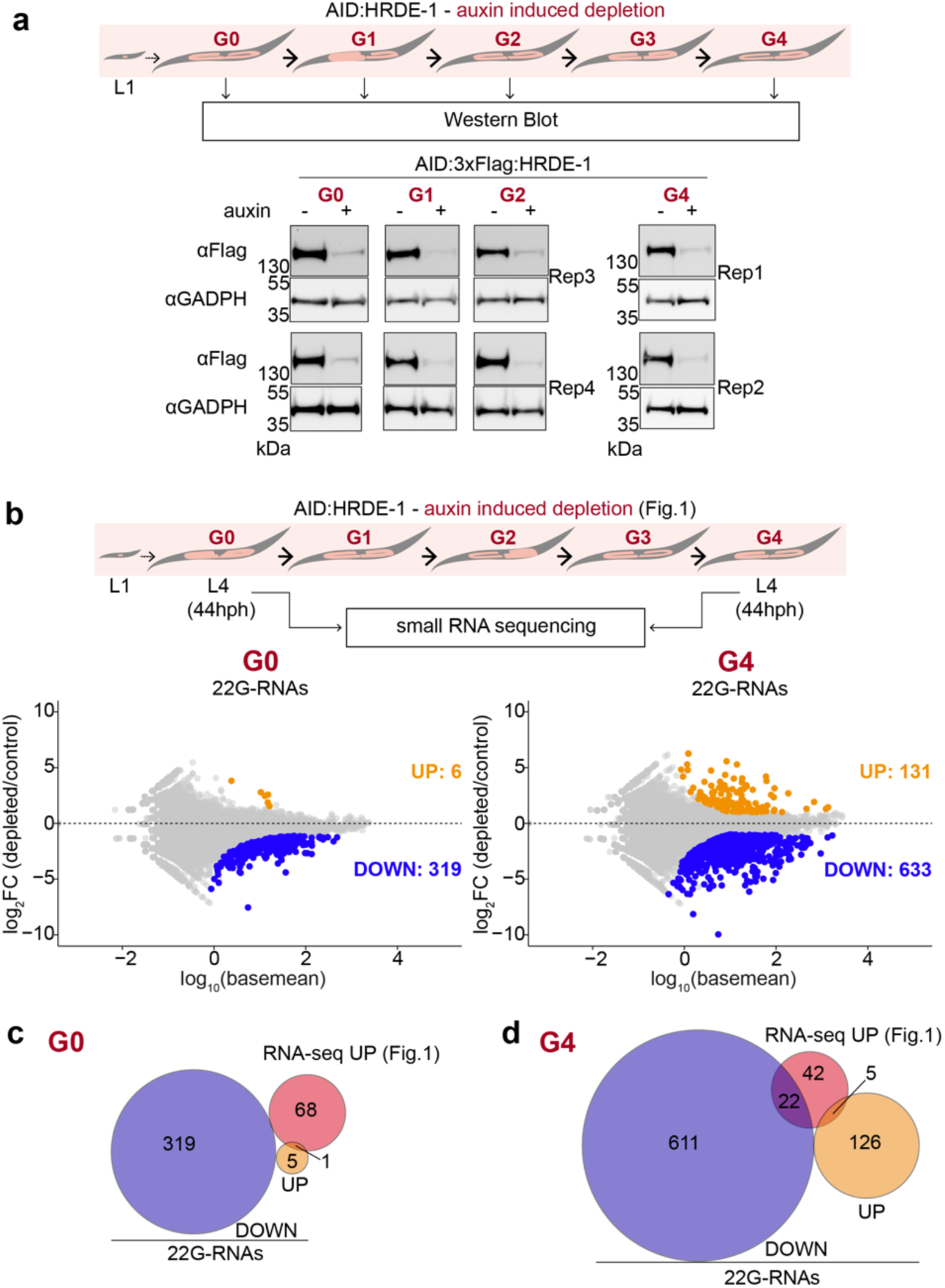
Transgenerational erosion of 22G-RNAs upon HRDE-1 depletion. **a**, Schematic of worm cultivation strategy for small RNA analysis and western blot confirming HRDE-1 depletion in sorted L4 larvae used for **Fig. 4b,d,e**. **b,** MA plot for 22G-RNA levels in G0 and G4 worms (same samples as in **Fig. 1b**). Orange, increased 22G-RNAs (log₂FC ≥ 1, FDR ≤ 0.05), blue, decreased 22G-RNAs (log₂FC ≤ −1, FDR ≤ 0.05). **c, d,** Venn diagrams showing overlap between the 69 HRDE-1–repressed genes (Fig. 1b) and those with significant 22G-RNA changes in G0 **(c)** and G4 **(d)**.

**Extended Data Fig. 7:**
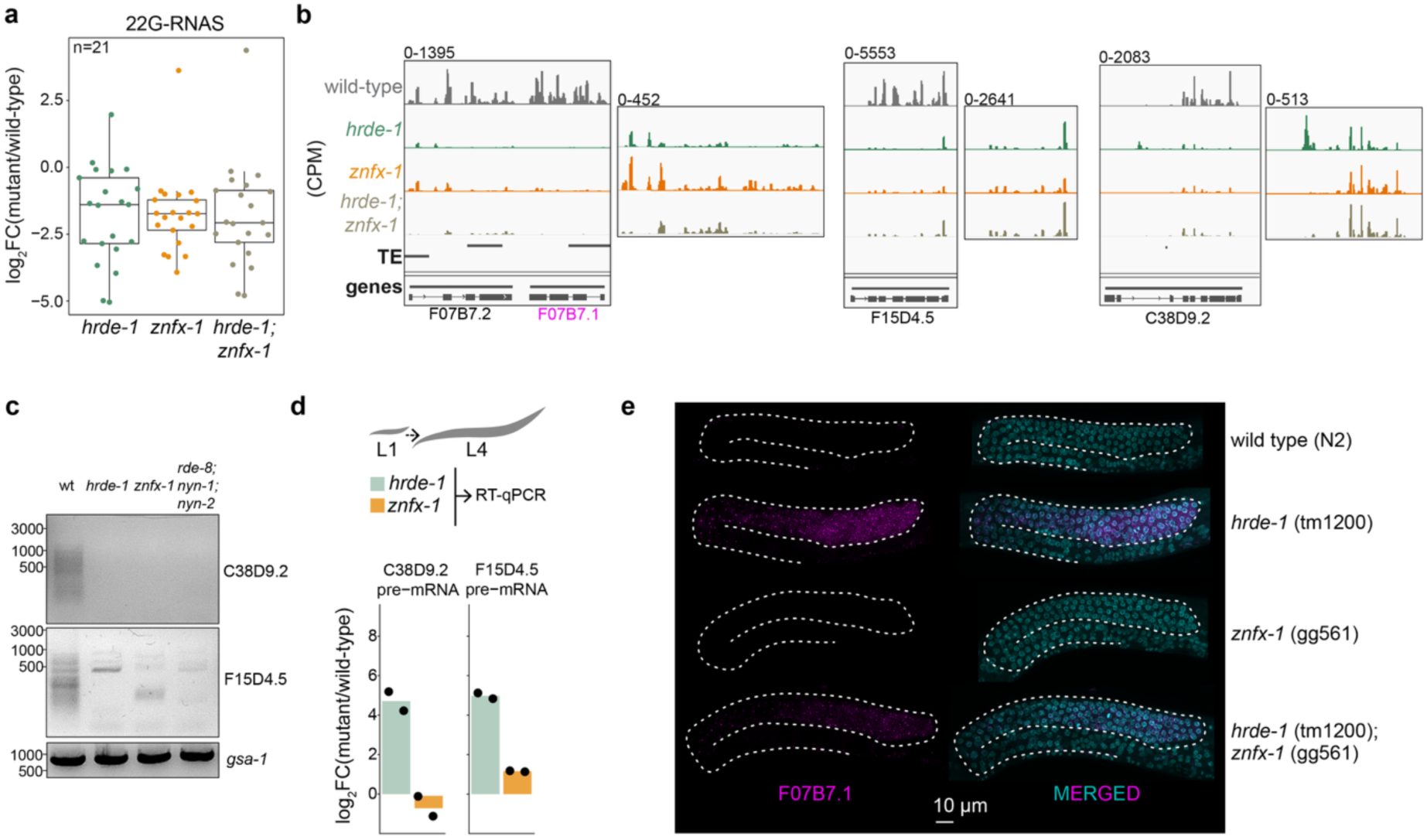
ZNFX-1 supports small RNA maintenance but is not required for transcriptional silencing of HRDE-1–regulated genes. **a,** Box plot of log₂ fold change (FC) of 22G-RNAs in *hrde-1*, *znfx-1*, and *znfx-1;hrde-1* mutants relative to wild-type for the 21 HRDE-1–repressed genes that lose small RNAs in G4. Data from Ouyang et al., 2022^14^. **b,** Genome browser views for representative targets (F07B7.2, F07B7.1 and F15D4.5) showing 22G-RNAs in wild-type (grey) *hrde-1* (green), *znfx-1* (orange), and *znfx-1;hrde-1* (beige). TE, mapped transposable elements. **c,** PolyUGylation assay of selected HRDE-1–repressed genes in wild-type (wt), *hrde-1*, *znfx-1*, and “endo” (*rde-8;nyn-1;nyn-2* triple mutant) backgrounds. *gsa-1* serves as a control. **d,** RT–qPCR of pre-mRNAs for F15D4.5 and C38D9.2 in *hrde-1* and *znfx-1* mutants at the L4 stage, normalized to *act-3*. Bars show mean log₂FC; dots represent biological replicates (*n* = 2). **e,** RNA FISH for *F07B7.1* (magenta) in wild-type, *hrde-1*, *znfx-1*, and *hrde-1;znfx-1* L4 larvae. DAPI (cyan) marks nuclei; gonads are outlined in dashed white.

**Extended Data Fig. 8.**
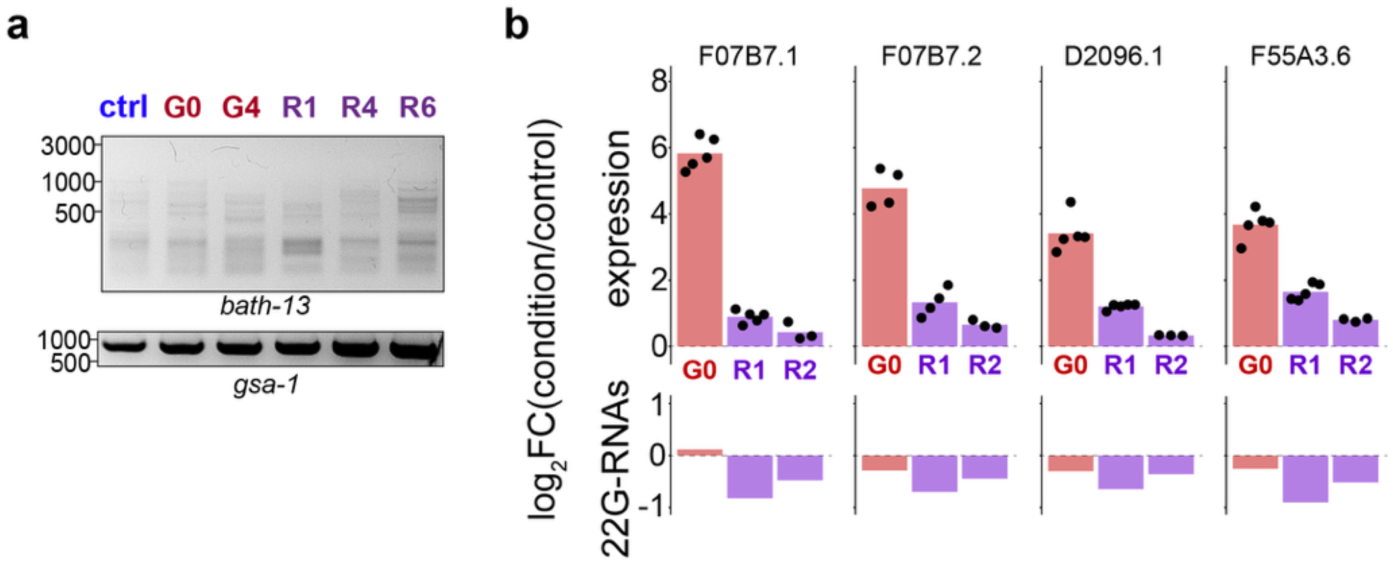
HRDE-1–dependent silencing recovery requires 22G-RNA biogenesis. **a,** PolyUGylation assay for *bath-13* across control (blue), G0 and G4 HRDE-1 depletion (red), and R1, R4, and R6 recovery generations (purple). *gsa-1* serves as control. **b,** RT–qPCR analysis of selected HRDE-1–repressed genes (as in **Fig. 6g**) in G0, R1, and R2 L4 worms, normalized to *act-3*. Bars show mean log_2_FC; dots represent biological replicates (*n* = 3–5). Bottom: corresponding average 22G-RNA fold change across generations.

## Methods

### Strains and worm culture

Worms were grown at 20°C in NGM plates and fed on *Escherichia coli* OP50 using standard methods^37^. For staged L4 larvae, worms were synchronized by bleaching. Briefly, gravid adults were incubated 5 min at 60 RPM in a 0.2 N NaOH and 1.5% NaOCl solution, washed three times, and incubated overnight for hatching with M9 buffer. 40,000 starved L1 larvae were placed on 150 mm NGM plates or NGM supplemented with Ethanol (0.2% v/v) or Auxin (indole-3-acetic-acid, 500 µM in Ethanol) for Auxin-inducible degradation experiments. For transgenerational experiments, L4 larvae were collected at each generation (44 hours post-hatching). Additionally, one plate was kept until ∼72 hph for bleaching gravid adults to obtain the next generation. The list of strains generated and used in this study is provided in **Supplementary Table 1**.

### Generation of CRISPR–Cas9 lines

Cas9-guide RNA (gRNA) ribonucleoprotein complexes were microinjected into the hermaphrodite syncytial gonad^38^. AID CRISPR-Cas9 tags were generated using double-stranded DNA repair templates (30 bp homology arms on each side) amplified by PCR using an AID::3xFlag or an AID:2xHA gBlock (IDT) sequence. Silent mutations were included, when needed, in the repair templates to prevent Cas9 cleavage. Mix concentrations were adapted from a previous study^39^. In brief, 10 µl mixes typically contained the following final concentrations: 0.1µg/µL Cas9-NLS protein (TrueCut V2, Invitrogen), 100 ng/µl in vitro transcribed target-gene gRNA, 80ng/µl of target-gene single-stranded DNA (ssDNA) repair template or 300 ng/μl target-gene double-stranded DNA (dsDNA) repair template, and 80 ng/µL pRF4 (roller marker). Cas9 and the target-gene gRNA were pre-incubated for 10-15 minutes at 37°C before adding the other components to the mixture. dsDNA repair templates were subjected to a melting/annealing step before addition to the final mix. A list of guide RNAs and primers used for CRISPR-Cas 9 lines is provided in **Supplementary Table 2**.

### Worm population sorting

Worms were prepared and collected as described before^6^. Briefly, we obtain large populations of precisely developmentally staged worms for genome-wide approaches, using a COPAS Biosorter as previously described^6^. For obtaining synchronous L4 population, among 40,000 to 80,000 larvae were harvested at 44 hph. To establish the sorting gate, for each sample, ∼ 100 worms were collected and inspected under a x40 objective brightfield microscope. The number of worms on each substage of the L4 development window was scored according to the vulva shape^40^. Larvae were sorted using a gate enriched for L4.4-4.5 larvae. All conditions of the same biological replicate were collected on the same day. 1,000 worm aliquots were harvested and spun down for 0.5 min at 0.7 g to remove sorting liquid, and flash-frozen in dry ice.

### Western Blot

Western blots were performed to monitor AID::3xFlag::HRDE-1 levels and its depletion upon auxin exposure; control worms were collected in parallel. For sorted larvae, 1,000 worms were resuspended in 50 µl of 4X NuPage LDS and 1X NuPage Reducing agent (Thermo Scientific™). For bulk larvae samples, 5 to 10 µl of packed worms were taken before extraction and mixed with 100 µl of 4X LDS and 1X Reducing agent. Worms were lysed by incubation at 95°C for 10 minutes. The sample was then centrifuged for 10 minutes at 20°C and 12,000 g, with ∼30-50% of the sample used for SDS-PAGE electrophoresis in 4-12% NuPage polyacrylamide gels. Gels were run in 1X MOPS buffer (Thermo Scientific™) and transferred to a Nylon membrane. Transfers were monitored with Ponceau staining and unstained before blocking. Primary antibody incubation was performed overnight at 4 °C with agitation. After, membranes were washed four times before secondary antibody incubation for 2 hours at room temperature and agitation. Antibody dilutions used: mouse anti-Flag 1:5,000 (Sigma-Aldrich, F3165), mouse anti-GADPH (Abcam, AB125247) 1:2,000, and Goat anti-mouse IgG HRP conjugated (Invitrogen, 31430) was used as secondary antibody, at a dilution of 1:10,000. The SuperSignal West Pico PLUS Chemiluminescent Substrate (Thermo Scientific) was used to detect the signal using a ChemiDoc MP imaging system (Biorad). Images are provided in **Source Data.**

#### RNA extraction

1 mL of TRIzol reagent (Invitrogen) was used following the manufacturer’s instructions. When extracting RNA from whole worms, 1,000-2,000 L4 sorted larvae or 10 µl of L4 (44 hph) packed worms were mixed with TRIzol and subjected to four to five cycles of freeze-thaw. Extraction was performed following the manufacturer’s instructions. 2 µl of GlycoBlue (15 mg/mL, Invitrogen) was added during the RNA precipitation step. Concentration and quality were estimated using Nanodrop (Thermo Scientific).

#### RT-qPCR

500 to 1,000 ng of total RNA was treated with 1 unit of Turbo DNAse (Invitrogen) for 30 min at 37°C and immediately purified using 1.8X solid-phase reversible immobilization beads for RNA purification (SpeedBeads Magnetic Carboxylase, Cytiva) and 3X Isopropanol. Reverse transcription was performed using M-MLV reverse transcriptase (Invitrogen) and random hexamers (Thermo Scientific). Expression was assessed by qPCR using Applied Biosystems PowerUp SYBR Green PCR Master mix and QuantStudio 3 Real-Time PCR System (Applied Biosystems). Three to five biological replicates were used, and expression changes were calculated to wild-type or non-treated samples, as indicated in the Figure legends. The primers used are listed in **Supplementary Table 3**.

#### PolyUGylated RNA detection

Gene-specific polyUG RNAs were detected by polyUG PCR as previously described^5^ (see also schematic in **Fig. 5b**). Briefly, 5 μg of total RNA was reverse-transcribed using M-MLV reverse transcriptase (Invitrogen) and 1 pg of RT primer. 1 μL of cDNA was used in a first PCR (20 cycles) using DreamTaq DNA polymerase (ThermoFisher). Next, PCRs were diluted at 1:100, and 1μL was used for a second PCR (30 cycles). Primers used are listed in **Supplementary Table 3**. Products were analyzed by agarose (2-3%) gel electrophoresis. Images are provided in **Source Data**.

### RNA fluorescent in situ hybridization (RNA FISH)

RNA FISH was performed as described before^6^. For *F07B7.1*, probes were designed in the exon regions that do not overlap with annotated transposable element sequences. We used unlabeled primary probes and Cy-5 fluorescently labeled secondary detector oligonucleotides (see **Supplementary Table 3**). Worms were harvested at 44 hph, incubated in fixative solution (3.7% Formaldehyde, 1X PBS) for 40 min at room temperature, and stored in 70% Ethanol. Fixated larvae were washed once in wash buffer (10% formamide, 2× SSC buffer) and hybridized with the corresponding FLAP-containing probes in 100 µL hybridization buffer (10% dextran sulfate, 2 mM vanadyl-ribonucleoside complex, 0.02% RNAse-free BSA, 50 µg E. coli tRNA, 2× SSC, 10% formamide) at 30 °C overnight. Hybridized larvae were washed twice with wash buffer, DAPI (7.5 ng/mL) was added in the second wash for nuclei staining and washed once in 2× SSC. For imaging, larvae were resuspended in 100 µL antifade buffer (0.4% glucose, 10 µM Tris-HCl pH 8, 2× SSC) with 1 µL catalase (Sigma-Aldrich) and 1 µL glucose oxidase (3.7 mg/mL, Sigma-Aldrich).

### Microscopy

Imaging was performed using an oil immersion x60, 1.42 numerical aperture objective on an inverted Andor BC43 spinning disk confocal microscope (Oxford Instruments). For image acquisition, the FUSION software was used, using the blue (405 nm) and red (638 nm) channels for DAPI and Cy-5 signals, respectively. Z-stacks were acquired with a 0.2 µM step size and a frame average of 2. Images were deconvoluted using the parameter “balanced”. Further image processing was done in FIJI^41^. Images were subtracted for background (rolling ball radius 50.0 pixels), maximal projections were obtained, and intensities were adjusted and applied to all the images corresponding to the same experiment. Uncropped images are provided in **Source Data**.

#### Nuclei isolation

For Antibody test experiments, 10,000 L4 larvae (wild-type and *met-2;set-25* strains) were obtained using a COPAS Biosorter as described above. We used this approach, since the *met-2;set-25* strain has an asynchronous development phenotype^42^. Worms were resuspended in 1 mL of cold Nuclear Extraction Buffer (NEB, 20 mM HEPES–KOH, pH 7.9, 10 mM KCl, 0.1% Triton X-100, 20% Glycerol, 0.5 mM spermidine)^18^ with 1X EDTA-free Protease Inhibitors (Thermo Scientific, 78425) and lysed by applying 40-50 strokes with a stainless-steel tissue grinder.

For germline nuclei isolation, 260,000 starved L1 larvae were plated in NGM-ethanol (control) or NGM-auxin plates (HRDE-1 or SET-25 germline-depletion) and grown for 44 h for L4 larvae harvesting. Worms were washed out of the plates and rinsed three times with 1X M9 buffer. Worm pellets were resuspended in 1.5 mL of cold NEB with 1X EDTA-free Protease Inhibitors and lysed by applying 65-70 strokes with a stainless-steel tissue grinder.

Lysis was monitored by inspection under a stereoscope (Zeiss), and the presence of GFP-positive nuclei was determined by mixing 2 µl of lysate with 2 µl of VECTASHIELD® Antifade Mounting Medium containing DAPI (Vector Laboratories) and inspection under a Zeiss Axio Imager M2 microscope equipped with a Princeton Instrument PIXIS 926 1024 camera, using Plan-Apochromat x20, 1.40 numerical aperture objective. Lysates were cleared by 4-minute centrifugation at 100 g and 4°C. Supernatant was transferred to a new tube and centrifuged for 4 minutes at 1,000 g and 4 °C to pellet the nuclei. Nuclei were resuspended in NEB and 1X EDTA-free Protease Inhibitors and used immediately for CUT&Tag (Antibody test experiments) or resuspended in NEB without glycerol and 1X EDTA-free Protease Inhibitors and kept on ice until sorting.

#### Germline nuclei sorting

Nuclei suspension was filtered and diluted 1:3-1:5 in cold NEB without glycerol and 1X EDTA-free Protease Inhibitors (Thermo Scientific™, 78425). Nuclei were analyzed and sorted using an Aria FACS III system (BD Biosciences). Aliquots of 500,000 GFP-positive nuclei were collected and kept on ice. After sorting, nuclei were concentrated by centrifugation for 4 minutes at 1,100 g and 4°C twice. Nuclei were resuspended in 100 µl of NEB and 1X EDTA-free Protease Inhibitors and stored at −20°C until further processing. For RNA extraction, nuclei were resuspended in 50 µl per 500,000 initial nuclei, and 1 mL of TRIzol LS (Invitrogen) was added before freezing. For GRO-seq, nuclei were resuspended in 30 µL of GRO-seq Nuclear freezing buffer^18^ and nuclear run-on was performed immediately, as described below.

#### Strand-specific RNA-seq library preparation

Total RNA was extracted and DNase-treated as detailed above. The DNase treatment step was skipped for RNA extracted from isolated germline nuclei. RNA integrity number (RIN) was assessed with the Agilent 2200 TapeStation system and used to determine the fragmentation time during library preparation. 100 ng of total RNA from whole worm or RNA from 500,000 germline nuclei were subjected to ribosomal RNA depletion using DNA-probed and DNase H as described before^20^. DNA probes were degraded by Turbo DNase treatment and RNA was purified using beads as explained before. Strand-specific libraries were prepared with NEBNext® Ultra™ II Directional RNA Library Prep Kit (E7770).

#### Small RNA library preparation

Libraries were prepared as described before^20^. Briefly, around 5 µg of total RNA was size separated in 15% NuPage 7% Urea polyacrylamide gels (Invitrogen) in 1X TBE buffer for 45 min at 180V. The small RNA fraction (between ∼17 to ∼25 nt) was extracted from the gel by incubation in 0.3 M NaCl and precipitated with 2 µL of GlycoBlue (Invitrogen) and 3 volumes of 100% Ethanol. After incubation on ice, the samples were centrifuged and washed once with 75% Ethanol. Isolated small RNAs were treated with 5’ Polyphosphatase (Biosearch Technologies), ligated with 3’ and 5’ end adaptors, and amplified by PCR using Illumina indexed primers and the NEBNext Ultra II Q5 Master Mix 2x (New England Biolabs). PCRs were purified with solid-phase reversible immobilization beads for DNA purification (SpeedBeads Magnetic Carboxylase, Cytiva) and analyzed using Agilent 2200 TapeStation System with high-sensitivity D1000 screentapes. Libraries were pooled and size selected using Pippinprep. Libraries were purified again using beads for DNA purification and quantified with the Qubit Fluorometer High Sensitivity dsDNA assay kit (Thermo Fisher Scientific, Q32851), based on a library size of 148 bp. An equimolar proportion of each library was mixed for sequencing.

#### GRO-seq library preparation

GRO-seq was performed as previously described^18^. Briefly, nuclear Run-On reactions were performed by incorporating 1 mM Bio-11-UTP (Lumiprobe), followed by RNA extraction with TRIzol and RNA fragmentation. Biotinylated nascent RNA was then purified using Dynabead^TM^ MyOne^TM^ Streptavidin C1 (Invitrogen). To prepare libraries, purified RNAs were first treated with T4 Polynucleotide Kinase (New England Biolabs) to repair the 5’-OH ends. This was followed by 3’ and 5’ adaptor ligation. Adaptor-ligated RNAs were then reverse transcribed using SuperScript IV Reverse Transcriptase (Thermo Fisher Scientific), following the manufacturer’s protocol with slight modifications: the reaction was incubated for 1 hour at 50°C and 10 minutes at 80°C. cDNA was PCR amplified using specific primers and the NEBNext Ultra II Q5 Master Mix 2x (New England Biolabs). Libraries were analyzed using the Agilent 2200 TapeStation System with high-sensitivity D1000 screentapes and quantified with the Qubit Fluorometer High Sensitivity dsDNA assay kit (Thermo Fisher Scientific, Q32851).

### CUT&Tag library preparation

Cut&Tag was performed following the EpiCypher protocol with a few modifications. Briefly. 100 μL of the nuclei resuspension generated as described above was used for binding to Concanavalin A beads (Epicypher 21-1401). Bead-bound nuclei were resuspended in 100 μL of Antibody150 buffer (20 mM HEPES pH 7.5, 150 mM NaCl, 0.01% digitonin, 2 mM EDTA, protease inhibitors, 0.5 mM spermidine), and 1 μg of primary antibody was added. 1:500 dilution of SNAP-CUTANA™ K-MetStat Panel (Epicypher) was added in this step in some experiments as an internal reference (spike-in) and for Antibody specificity assessment, as indicated in the text. Reactions were incubated overnight in a thermocycler shaker set up at 4°C with shaking cycles at 2000 rpm, 15 seconds ON 45 seconds OFF. 1 μg of secondary antibody in 100 μL of Digitonin150 buffer was added (20 mM HEPES pH 7.5, 150 mM NaCl, 0.01% digitonin, protease inhibitors, 0.5 mM spermidine), followed by tagmentation with CUTANA pAG-Tn5 (Epicypher 15-1117) in tagmentation buffer (20 mM HEPES pH 7.5, 300 mM NaCl, 0.01% digitonin, protease inhibitors, 0.5 mM spermidine, 10 mM MgCl_2_). Resulting libraries were amplified with i5 and i7 primers using 13 to 15 PCR cycles and purified using beads for DNA purification (SpeedBeads Magnetic Carboxylase, Cytiva). Libraries were analyzed using the Agilent 2200 TapeStation System with high-sensitivity D1000 screentapes and quantified with the Qubit Fluorometer High Sensitivity dsDNA assay kit (Thermo Fisher Scientific, Q32851). Three or four biological replicates were performed for each experiment. For the antibody test experiments, we used the following anti-H3K9me3 antibodies: Abcam (ab8898), Diagenode (C15410193), Cell Signaling (1396), and Active Motif (AM39161), which were selected for further experiments (see **Extended Data Figure 3**). Additionally, we used the anti-H3K4me3 (Active Motif, AM61379) and anti-H3K23me3 (Active Motif, AM61499) primary antibodies. We used Guinea pig anti-Rabbit (Antibodies-Online, ABIN101961) or Rabbit anti-mouse (Abcam, ab46540) IgG, according to the source of the primary antibody.

Library sequencing.

Multiplexed libraries were sequenced on NextSeq500 or NextSeq2000 Illumina platforms.

### Sequencing data analysis

All the sequencing data were demultiplexed with Illumina bcl2fastq converter (version v2.17.1.14), and quality control was performed with fastQC (version v0.11.5).

#### RNA-seq analysis

RNA-seq reads were aligned to the WBcel235 genome using STAR^43^ and quantified by RSEM in parallel^44^ using the RSEM-STAR pipeline with additional options ‘--calc-pme --calc-ci --estimate-rspd’. Further analysis was conducted using the Bioconductor DESeq2 package^45^ from gene-level values for rounded RSEM counts. Genes with at least 10 normalized counts in all replicates were considered for differential expression analysis. Differential expressions between replicate groups were assessed using the Wald Test, thresholding for significance at FDR ≤ 0.05.

#### Small RNA-seq analysis

The 3’ adaptor (TGGAATTCTCGGGTGCCAAGG) was trimmed from the raw reads using cutadapt (version 4.9). The 5’ and 3’ 4 nt UMIs were removed using cutadapt (version 4.9) with options -u 4 and -u −4. After removing UMIs, the reads from 18 to 24 nt were selected using bioawk. The size-selected reads were mapped on the *C. elegans* genome (WBcel235) using bowtie2 (version 2.5.4) with options -L 6 -i S,1,0.8 -N 0.

Mapped reads were used to estimate the abundance of small RNAs derived from structural RNAs using subread/2.0.0 featureCounts with options -O -M --fraction and annotations corresponding to tRNA, snRNA, snoRNA, rRNA or RNA (as annotated in the iGenome distribution of WBcel235 obtained at ftp://igenome:G3nom3s4u@ussd-ftp.illumina.com/Caenorhabditis_elegans/Ensembl/WBcel235/Caenorhabditis_elegans_Ensembl_WBcel235.tar.gz). The abundance of non-structural RNAs was estimated by subtracting the above counts from the number of mapped reads. Initially mapped reads were classified using a custom Python program according to their length, composition, and the annotations on which they mapped. Reads that did not match miRNA and piRNA annotations were considered as potential endo-siRNAs. The potential endo-siRNAs of size 21 to 23 nt that started with G were classified as 22G-RNAs if they mapped antisense to annotation belonging to the following categories: DNA transposons, RNA transposons, satellites, simple repeats (as annotated in http://hgdownload.cse.ucsc.edu/goldenPath/ce11/database/rmsk.txt.gz) or pseudogene or protein-coding genes (as annotated in the iGenome distribution of WBcel235 obtained at ftp://igenome:G3nom3s4u@ussd-ftp.illumina.com/Caenorhabditis_elegans/Ensembl/WBcel235/Caenorhabditis_elegans_Ensembl_WBcel235.tar.gz). The resulting alignment was used to generate the normalized bigwig file using millions of non-structural RNAs as normalizer. This was done using bedtools (version 2.31.1) bamCoverage. Genome_build: C. elegans ce11 (WBcel235).

Publicly available data, corresponding to the BioProject PRJNA819556^14^, was downloaded from the SRA repository and re-analyzed following the processing steps mentioned above, omitting the UMIs removal step.

#### GRO-seq analysis

Analysis of the sequencing data was performed as previously described^46^. The scripts and workflows are available at https://gitlab.pasteur.fr/bli/bioinfo_utils. Reads were normalized to millions of non-structured RNAs. Plots in **Fig. 2f,g**, were generated using deeptools (version 3.5.6)^47^, with plotProfile and plotHeatmap tools, respectively.

#### CUT&Tag

Reads were aligned with Bowtie 2^48^ to the WBcel235 genome, with options “--local --very-sensitive-local --dovetail --soft-clipped-unmapped-tlen”. Reads were additionally deduplicated using samtools. Normalization was calculated by counts multiplied by one million divided by the total number of reads. For normalization using the SNAP-CUTANA™ K-MetStat Panel (Epicypher), spike-in abundance was quantified directly from raw FASTQ files before alignment. For each sample, both R1 and R2 FASTQ files were searched for exact matches to each spike-in sequence. Counts are summed to produce a per-sample total spike-in abundance. This abundance is converted to a normalization factor. All bigwig coverage tracks are scaled using this factor.

Peaks were called using MACS2^49^ with “callpeak -q 0.05 -g ce --keep-dup all”. Peaks were further filtered to have MACS2 FDR < 0.01 in merged reads from either wild-type or mutant, or non-treated or treated, replicates and FDR < 0.05 in at least two individual replicates. Quantifications for the selected set of peaks were done for each dataset using subread featureCounts^50^. Next, these quantifications were analyzed using a DESeq2 generalized linear model.

### Statistics and reproducibility

All the experiments shown in this study were performed independently at least twice, and the exact number of biological replicates is indicated in each figure legend. Statistical test and exact P values are shown in the figures and indicated in the corresponding legend. Other calculations and plots, in addition to sequencing data analysis explained above, were done in RStudio. No statistical methods were used to predetermine sample sizes. Gene lists used in this study are provided in **Supplementary Table 4**.

## Data availability

All the sequencing data are available at the following accession numbers: Whole worm RNA-seq GSE311522, Small RNA-seq GSE311555, nuclear RNA-seq GSE311553, nuclear GRO-seq GSE311554, Whole worm CUT&Tag GSE311918, and germline nuclei CUT&Tag GSE311921. Source Data are provided for this study.

## Acknowledgments

We would like to thank all the members of the Cecere laboratory for the helpful discussions on the manuscript and suggestions on the project. We thank Dr. Florian Muller, head of the Photonic BioImaging platform, and Dr. Gaëlle Letort, research engineer in the Department of Developmental and Stem Cell Biology, for their assistance on RNA FISH imaging interpretation. We thank the Caenorhabditis Genetics Center (CGC) for *C. elegans* strains, supported by the National Institutes of Health - Office of Research Infrastructure Programs (P40 OD010440). We thank Carolyn Phillips for sharing *rde-8* and *nyn-1;nyn-2* strains. We thank the Flow Cytometry, Photonic BioImaging, and Biomics facilities at Institut Pasteur. This project has received funding from the Institut Pasteur, the CNRS, and the European Union’s Horizon 2020 research and innovation program (grant agreement no. 101002999). A.M.L.R. received funding from the European Union under the HORIZON-MSCA-2022-PF-01 program no. 101109836 — MOMENTS and A.K.M. received funding from LabEx Revive fellowship ANR-10-LBX-73.

## Author contributions

A.M.L.R and G.C. identified and developed the core questions of the work and supervised the whole project, contributed to the data analysis, and wrote the manuscript. A.C. developed pipelines for small RNA-seq, RNA-seq, and CUT&Tag analysis, and assisted A.M.L.R. in downstream data examination. A.K.M. developed the germline nuclei isolation protocol, and together with J.J. and A.M.L.R., adapted it for downstream genomic approaches (total RNA-seq, CUT&Tag, and GRO-seq). P.Q. and A.M.L.R. performed antibody screenings for the CUT&Tag protocol. A.M.L.R. and J.J. performed CUT&Tag experiments. A.M.L.R. performed RNA-seq, small RNA-seq, GRO-seq, and RNA FISH experiments. L.B. performed together with O.F. the RNAi inheritance assay. A.M.L.R. and S.I. performed the polyUGylation assays. A.M.L.R. and S.C. performed RT-qPCR of AID and mutant strains. L.B. assisted A.M.L.R. with the COPAS Biosorter operation for staged-worm sorting. L.B. and J.O. generated strains and crosses used in this project and helped with worm growing and maintenance.

## Materials & Correspondence

Correspondence and material requests should be addressed to Germano Cecere

